# Acute and cumulative light exposure differentially influence mood in healthy adults

**DOI:** 10.64898/2026.07.31.742017

**Authors:** Chloe Roddis, Altug Didikoglu, Aghileh Ebrahimi, Amy L. Gillespie, Catherine J. Harmer, Beatriz Bano-Otalora, Nina Milosavljevic

## Abstract

Light influences mood in both clinical and experimental settings, yet it remains unclear how ambient light exposure relates to mood in daily life, which dimensions of mood are most sensitive to light and over what timescales these associations emerge. Here, we combined continuous personal light monitoring using wearable light sensors with repeated assessments of multiple dimensions of mood and affect, including self-reported mood ratings and cognitive measures of affective bias, alongside physiological monitoring in 49 healthy adults living under naturalistic conditions. Brighter light was consistently associated with more positive self-reported mood. Associations emerged over distinct temporal windows, with greater melanopic irradiance during the preceding 30 min associated with increased feelings of *energy*. Importantly, this association remained significant after accounting for alertness, physical activity and sleep-related covariates, suggesting that it is not solely explained by the established alerting effects of light. In contrast, several positive mood dimensions, including *happiness*, feeling *accompanied*, *enjoyment*, *motivation* and *relaxation*, were associated with light accumulated over longer periods (60-240 min). Furthermore, *happiness*, feeling *accompanied*, *enjoyment* and *relaxation,* but not *energy,* were also associated with greater cumulative daily exposure to light above 250 lux melanopic equivalent daylight illuminance (EDI), a threshold proposed to support healthy daytime light exposure. Behavioural measures of affective bias showed no robust associations with light exposure. Together, these findings indicate that everyday light exposure is associated with distinct dimensions of mood over different timescales, highlighting a complex relationship between light and affect in daily life.

**Significance Statement:** Light is a promising target for improving mental health, yet little is known about how everyday light exposure relates to mood outside laboratory and clinical settings. Using wearable light sensors and repeated mood assessments in healthy adults living under naturalistic conditions, we show that different dimensions of mood are linked to light exposure over distinct timescales, from recent exposure to cumulative light history. Importantly, the association between recent light exposure and feelings of *energy* remained independent of physical activity and sleep-related factors. These findings demonstrate that effects of light on mood are dynamic and multidimensional, highlighting the need to move beyond simple measures of total light exposure and informing future efforts to design healthier light environments that support emotional wellbeing.

## Introduction

Ambient light is a primary regulator of daily rhythms in human behaviour and physiology, influencing sleep-wake cycles, alertness, mood regulation and cognitive function (1–3). However, modern patterns of light exposure have diverged markedly from natural light-dark cycles, characterised by reduced daytime light exposure and increased exposure to artificial light at night (4–8). This has raised important questions about whether insufficient or mistimed light may contribute to mood and affective vulnerability. Consistent with this possibility, light is an established therapeutic intervention for seasonal and non-seasonal depression, demonstrating that appropriately timed light exposure can causally influence mood symptoms (9–14). Large-scale population studies also suggest that greater daylight exposure is associated with lower risk of depression and better mood-related outcomes (4, 5). Nevertheless, despite substantial evidence linking light exposure to mood from clinical, epidemiological and controlled laboratory studies, far less is known about how everyday light exposure influences momentary mood in healthy individuals during daily life.

A key challenge in characterising these relationships is that the biological effects of light are inherently multidimensional. Light exposure varies not only in total amount, but also in its timing, intensity, duration, temporal pattern and spectral composition, each of which can independently influence its physiological and affective relevance (15–17). Although these dimensions are known to modulate multiple aspects of circadian physiology, including circadian entrainment, phase shifting, melatonin suppression, sleep-wake regulation, alertness and circadian amplitude, their influence on affective state remains less well understood (18–32). Recent advances in wearable light-sensing technologies, including wrist-worn actigraphy and smartphone-based assessments, have enabled continuous, high-resolution monitoring of personal light exposure in daily life, revealing associations between habitual light environments and sleep timing, sleepiness, alertness, cognitive performance and depressive symptoms (6, 33-38).

Cumulative findings across the field support the use of melanopic equivalent daylight illuminance (melanopic EDI), a metric that quantifies the non-visual effects of light mediated by intrinsically photosensitive retinal ganglion cells (ipRGCs), particularly with respect to circadian entrainment, sleepiness and alertness (39, 40). Given the established links between circadian phase and affect, with subjective happiness and cheerfulness varying across the circadian cycle (41), light-induced changes in circadian timing may influence mood through downstream circadian mechanisms. In addition to these indirect circadian and sleep-mediated pathways, light can also exert more direct effects on alertness, arousal and emotional processing, suggesting that mood-relevant effects depend not only on overall light exposure but also on the timing and timescale over which exposure is assessed (2, 21, 30, 42–50).

Importantly, mood is multidimensional and can be assessed through both subjective self-report measures and objective indices of emotional processing. One such index is affective bias, reflecting the preferential processing of positive versus negative emotional information. This is altered across several psychiatric disorders and is considered a clinically relevant transdiagnostic process and potential treatment target (51, 52). Performance-based measures of affective bias may therefore complement self-reported outcomes, by capturing affective function, while also being less susceptible to demand characteristics and treatment expectancy effects (53).

Despite growing evidence linking light exposure to affective functioning, two key questions remain unanswered: which dimensions of mood and affect are most sensitive to light exposure, and over what timescale these effects emerge, ranging from acute exposure in the minutes preceding assessment to cumulative light exposure across the day. Given the multidimensional nature of affect and the multiple pathways through which light may influence mood, we hypothesised that acute and cumulative light exposure would show distinct associations with different dimensions of subjective mood and affective bias in everyday life. To test this, we characterised individual profiles of melanopic EDI and examined their relationship with affect in individuals with no mental health diagnoses under naturalistic conditions. In a one-week observational study, we continuously monitored personal light exposure using wearable ActLumus devices alongside physiological activity measured with Fitbit trackers. Participants also completed repeated smartphone-based mood assessments throughout the day, a daily behavioural task measuring affective bias and a morning sleep diary. We found that even brief light exposures, occurring within as little as 30 minutes, were associated with increased self-reported feelings of *energy, happiness* and *motivation*. Interestingly, the association with feelings of *energy* remained significant after adjusting for covariates, including activity levels and sleep-related measures, whereas associations with *happiness* and *motivation* emerged only at longer durations of light exposure (60-180 min). More broadly, short-term and cumulative light exposure showed distinct associations with mood, consistent with effects on dynamic, state-like aspects of affect. In contrast, behavioural measures of affective bias, which may reflect more stable, trait-like processes, showed little evidence of association with light exposure.

## Results

### Longitudinal Light Exposure in Everyday Life

Light exposure data, in addition to self-reported sleep, were collected from 51 participants (49 after cleaning) throughout their everyday life for a 1-week period between December 2023 to September 2024. Sunrise and sunset times during the study period varied due to season, with sunrise hours ranging between 05:44 and 08:25 and sunset between 15:49 and 22:43. Wrist-worn sensors were selected for their practicality and participant compliance (6, 33), though they do not perfectly capture light at eye level and may occasionally be obscured by clothing (54). To mitigate this, participants were instructed to keep the devices uncovered whenever possible to optimise measurement accuracy. Participants were mostly under 30 years old (mean ± SD: 27.35 ± 7.03 years; 75% <30 years) with an approximately equal sex distribution (49% male, 51% female). The study sample included participants who are students (29%) or full-time workers (61%) (SI Appendix, Table S1). Demographic and mental health assessments confirmed that the recruited population did not include individuals with diagnosed depression or anxiety disorders (Table 1). Average 24-h light exposure was 4.06 lux melanopic EDI (SD = 31.13), with a mean daytime melanopic EDI of 31.19 lux between 07:00 and 20:00 (Fig. 1A-1B). Across records for light exposure, 37% of samples were on free days, compared to 63% on workdays. Light exposure was significantly higher on weekdays when averaging overall weekday exposure, compared to weekends (*t*(250,383) = -25.31, *p* < 0.001), with the largest differences observed in the morning (Fig. 1C). A nonlinear least-squares cosine model fitted to the pooled log-transformed melanopic EDI data identified a significant 24-hour rhythmic component (amplitude magnitude = 1.515, SE = 0.002, *p* < 0.001). The estimated baseline log melanopic EDI was 0.612 (SE = 0.001, *p* < 0.001). The model peak occurred at ∼14:23, corresponding to early afternoon (Fig. 1D). The mean daily timing of light exposure above 250 melanopic EDI lux occurred at 13:43 local time (interquartile range [IQR] 11:18-16:06, range 06:24-04:00 next day). Time spent over 250 lux melanopic EDI was zero on 5.2% of days and did not differ between weekdays and weekends (*t*(185.23) = -0.34, p = 0.73). Individuals spent on average 175 minutes over the recommended melanopic EDI for daytime hours from Brown et al. (2022), (>250 lux melanopic EDI) and 73 minutes over bright light (>1,000 lux melanopic EDI) (Fig. 1E-1F; Fig. S1A-B). It is important to note, however, that these recommendations were developed for corneal light exposure, whereas the present measurements were obtained from wrist-worn devices.

**Fig. 1.**
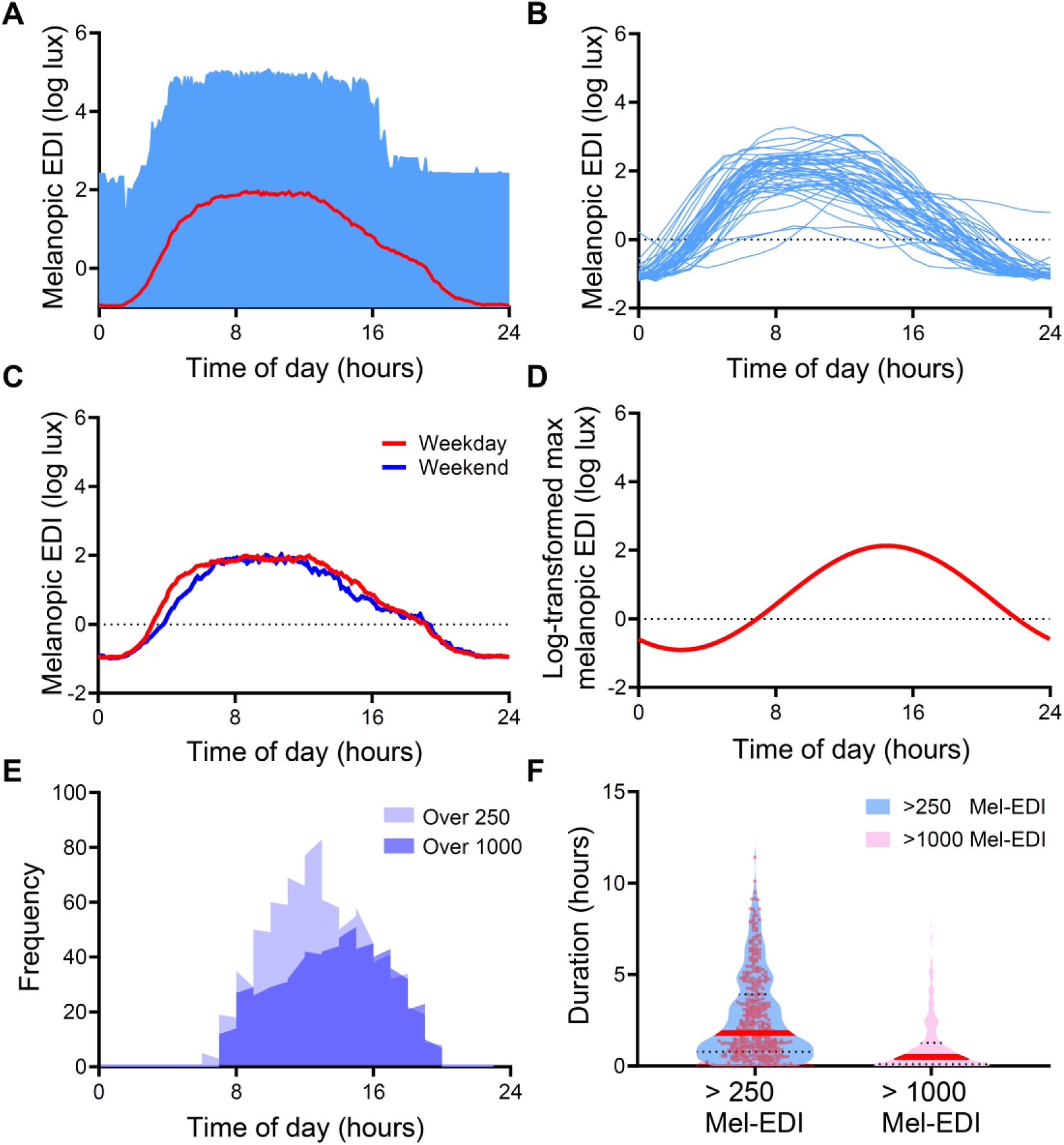
Daily patterns of light exposure across the 24-h day. Data are from 49 participants after data cleaning. (**A**) Mean melanopic equivalent daylight illuminance (melanopic EDI) across the day for all participants. The red line represents the mean log-transformed melanopic EDI, and the shaded ribbon indicates the minimum-to-maximum range. (**B**) Individual participants’ light-exposure profiles across the day; blue lines represent individual participants. (**C**) Comparison of mean melanopic EDI across the 24-h day between weekdays and weekends. (**D**) Cosine fit to the log-transformed maximum melanopic EDI across the day, aggregated across all participants. (**E**) Frequency distribution of melanopic EDI as a function of time of day. Cumulative exposure frequency is shown for melanopic EDI levels greater than 250 lx (light blue) and greater than 1,000 lx (dark blue). (**F**) Violin plots showing the duration of daytime exposure (08:00–20:00 hours) to melanopic EDI greater than 250 lx (blue) and greater than 1,000 lx (pink).

**Table 1.**
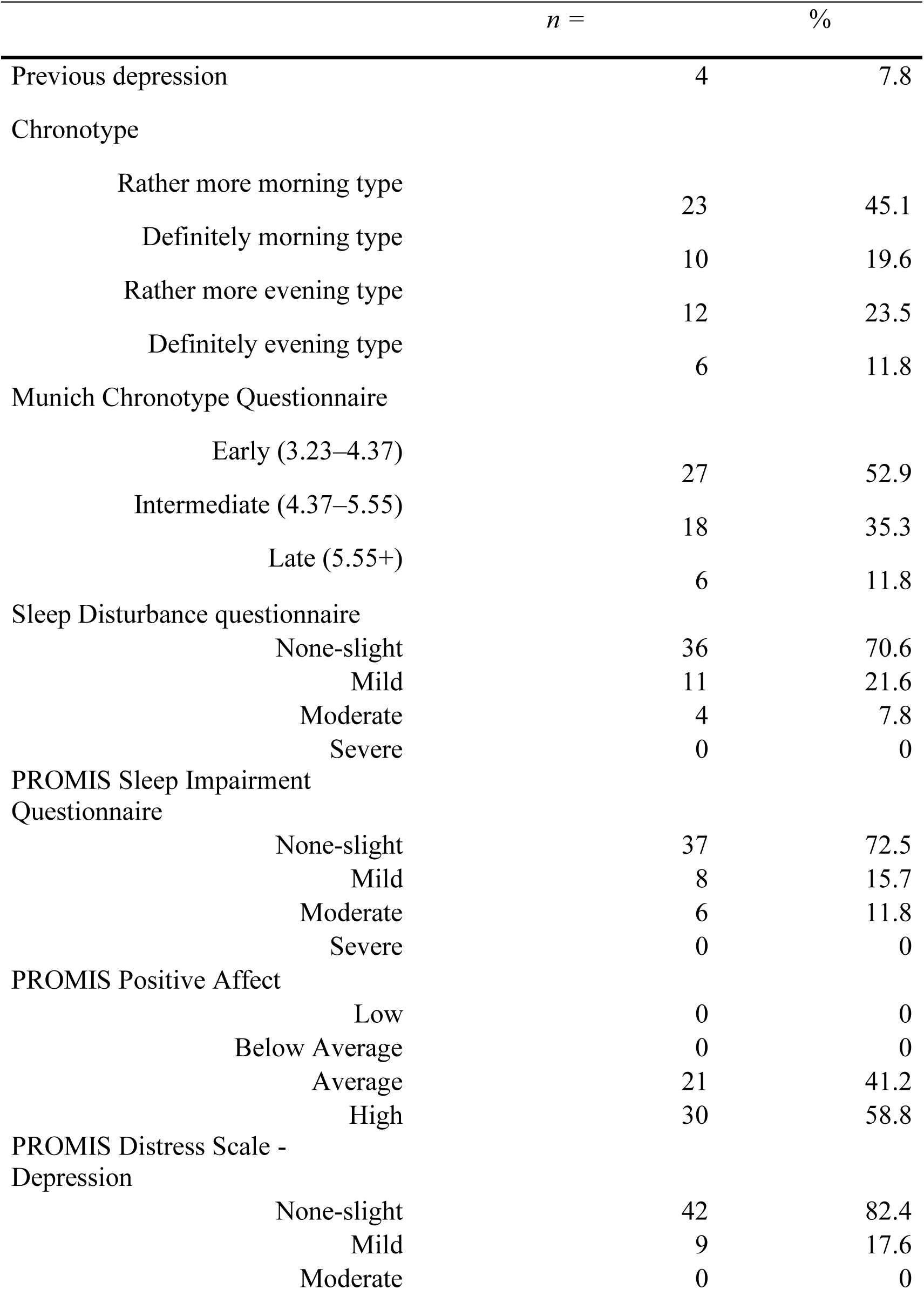
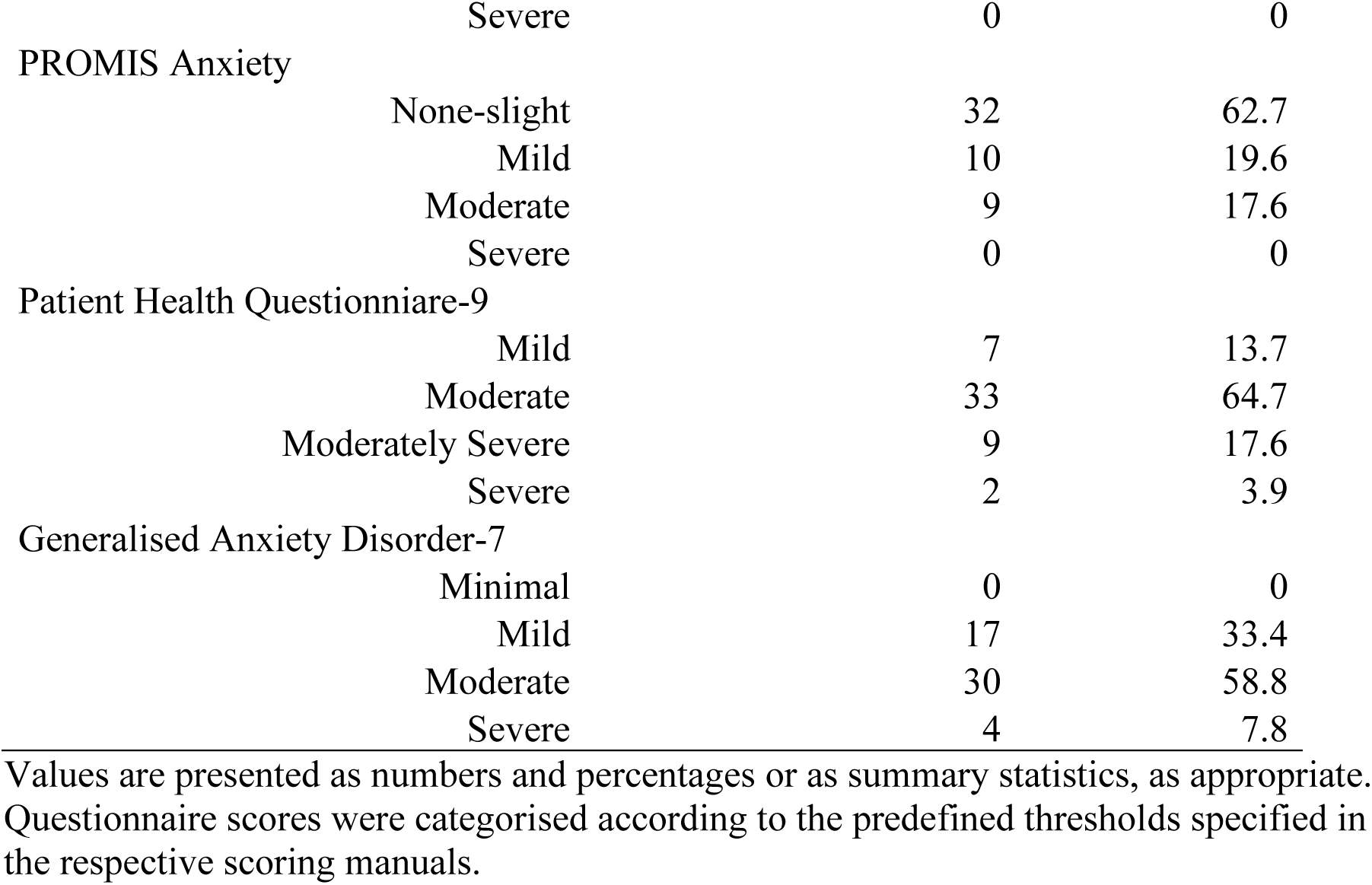
Demographic characteristics and baseline questionnaire scores of healthy participants (n = 51).

|  | *n* = | % |
| --- | --- | --- |
| Previous depression | 4 | 7.8 |
| Chronotype |  |  |
| Rather more morning type | 23 | 45.1 |
| Definitely morning type | 10 | 19.6 |
| Rather more evening type | 12 | 23.5 |
| Definitely evening type | 6 | 11.8 |
| Munich Chronotype Questionnaire |  |  |
| Early (3.23–4.37) | 27 | 52.9 |
| Intermediate (4.37–5.55) | 18 | 35.3 |
| Late (5.55+) | 6 | 11.8 |
| Sleep Disturbance questionnaire |  |  |
| None-slight | 36 | 70.6 |
| Mild | 11 | 21.6 |
| Moderate | 4 | 7.8 |
| Severe | 0 | 0 |
| PROMIS Sleep Impairment Questionnaire |  |  |
| None-slight | 37 | 72.5 |
| Mild | 8 | 15.7 |
| Moderate | 6 | 11.8 |
| Severe | 0 | 0 |
| PROMIS Positive Affect |  |  |
| Low | 0 | 0 |
| Below Average | 0 | 0 |
| Average | 21 | 41.2 |
| High | 30 | 58.8 |
| PROMIS Distress Scale - Depression |  |  |
| None-slight | 42 | 82.4 |
| Mild | 9 | 17.6 |
| Moderate | 0 | 0 |

|  |  |  |  |
| --- | --- | --- | --- |
|  | Severe | 0 | 0 |
| PROMIS Anxiety | None-slight | 32 | 62.7 |
|  | Mild | 10 | 19.6 |
|  | Moderate | 9 | 17.6 |
|  | Severe | 0 | 0 |
| Patient Health Questionnaire-9 | Mild | 7 | 13.7 |
|  | Moderate | 33 | 64.7 |
|  | Moderately Severe | 9 | 17.6 |
|  | Severe | 2 | 3.9 |
| Generalised Anxiety Disorder-7 | Minimal | 0 | 0 |
|  | Mild | 17 | 33.4 |
|  | Moderate | 30 | 58.8 |
|  | Severe | 4 | 7.8 |

Values are presented as numbers and percentages or as summary statistics, as appropriate.
Questionnaire scores were categorised according to the predefined thresholds specified in
the respective scoring manuals.

### Daily Assessments of Momentary Self-reported Mood

Participants were free to determine when they reported their momentary self-reported mood up to five times per day on a 0-100 scale, capturing seven dimensions: *accompanied, carefree, energetic, enjoyment, happy, motivated* and *relaxed*. After excluding invariant responses (all 0 or 100) and participants with >7 missing entries per week, 1,589 observations from 50 participants (after cleaning) across 407 days were analysed, with a median of 32 entries per participant. Across all self-reported mood variables, median scores were consistently above 50, suggesting generally positive mood (Fig. 2A). Reporting times spanned ∼05:30-03:00 (median 14:30), with entries occurring 0-20.5 h after waking and median alertness (KSS = 4), reducing concern about time-of-day bias (Fig. 2B, SI Appendix, Fig. S3A-B).

**Fig. 2.**
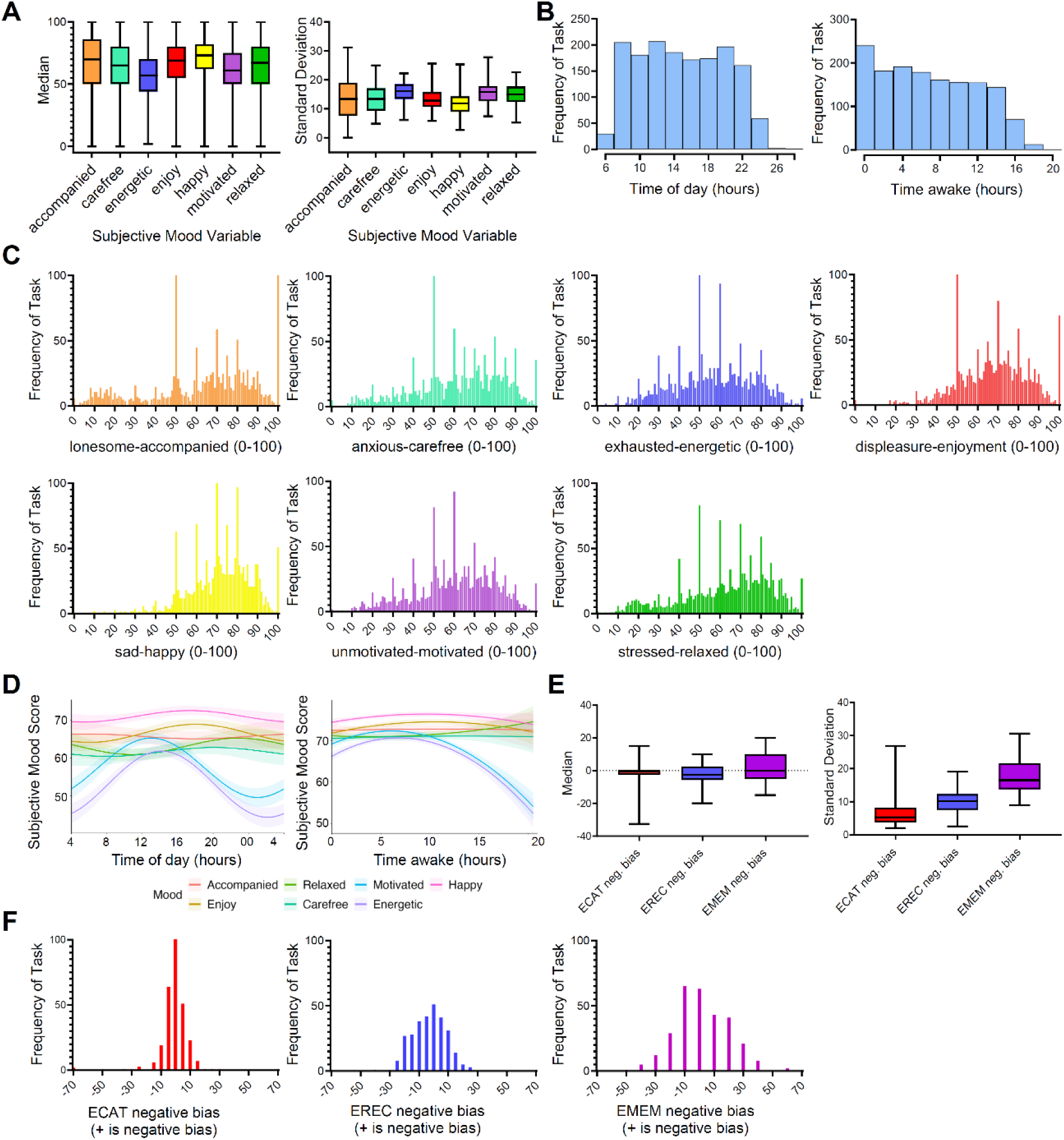
Variation in self-reported mood scores and behavioral affective bias scores across the week. Data from 50 participants were available for the self-reported mood measures and from 47 participants for the affective bias measures. For each mood rating, participant-level summary statistics were calculated, and cosine models were fitted to assess time-of-day effects. (**A**) Boxplots showing participant-level median values and within-person standard deviations for the self-reported mood scores. (**B**) Bar charts showing the frequency distribution of self-reported mood questionnaire completions across time of day and time awake. (**C**) Bar charts showing the frequency distribution of all self-reported mood ratings on a 0–100 scale. (**D**) Cosine-model fits used to assess circadian rhythmicity in self-reported mood, modeled as a function of time of day and time awake. Each line represents a different self-reported mood score, as indicated in the key. (**E**) Boxplots showing participant-level median values and within-person standard deviations for affective bias scores. Positive scores indicate a more negative bias, whereas lower scores indicate a more positive bias. Negative bias measures for emotional categorisation (ECAT), emotional recall (EREC), and emotional memory (EMEM) are shown as ECAT, EREC, and EMEM scores, with higher values indicating greater negative bias. (**F**) Bar charts showing the frequency distribution of behavioral affective bias scores across all participants. ECAT, emotional categorisation; EREC, emotional recall; EMEM, emotional memory.

Self-reported mood dimensions also differed in overall variability (Fig. 2C). Fluctuations in mood variables consistent with a state-like nature and sensitivity to contextual influences, displayed the largest fluctuations, including e*nergetic* (median = 57, SD = 19.33), *motivated* (median = 60, SD =19.52)*, relaxed* (median = 66, SD = 21.24), *carefree* (median = 64, SD = 21.87) and lastly *accompanied* (median = 70, SD = 26.45). In contrast, *happy* and *enjoyment* were comparatively stable (median = 73, SD = 15.59; median = 69, SD = 17.73), reflecting a more trait-like profile. Summary metrics (median, SD, amplitude, mesor, cosine residuals) confirmed these patterns: variability measures (SD, amplitude) were highest for *energetic* and lowest for *happy*, indicating individual differences in mood dynamics (Fig. 2A;SI Appendix, Fig. S2A-C).

Cosine models were applied to assess circadian rhythmicity in self-reported mood. Standardised coefficients and p-values showed that cosine was negatively associated with *energetic* (β = -0.24, *p* < 0.05) and *motivated* (β = -0.24, *p* < 0.05), indicating more *energetic* and *motivated* moods in the middle of the day (Fig. 2D). Whereas *happy, relaxed, carefree, enjoyment* and *accompanied* showed low amplitude variation or nonsignificant rhythmicity (SI Appendix, Table S2; Fig. S2A). Additionally, self-reported mood stays stable for the first 10-12 hours awake, after which motivation and energy decline (Fig. 2D). Amplitude and mesor estimates characterised rhythm-adjusted means and strength (SI Appendix, Fig. S2B-C).

Both individual-level factors and day-to-day variability significantly contributed to mood variance across all seven dimensions (*Table 2*). For an energetic mood, for example, effects of individual differences (*F*(50,1182) = 14.03, *p*<0.001, η² = 0.285), the day of assessment (*F*(10, 1182) = 4.81, *p* < 0.001, η² = 0.020) and their interaction (*F*(346, 1182) = 1.54, *p* < 0.001, η² = 0.216) were significant. This pattern was consistent across all self-reported mood variables (Table 2).

**Table 2.** Variance in outcome measures explained by participant and task day (n = 50)

| Emotion | Source | df | Sum of Squares | Mean Square | F Value | P value |
| --- | --- | --- | --- | --- | --- | --- |
| Accompanied | Task Day | 10 | 7,658 | 765.80 | 3.89 | <.001 *** |
|  | Subject | 50 | 719,611 | 14,392.22 | 73.16 | <.001 *** |
|  | Task Day x Subject | 346 | 150,978 | 436.32 | 2.22 | <.001 *** |
|  | Residuals | 1,182 | 232,539 | 196.68 |  |  |
| Carefree | Task Day | 10 | 6,080 | 608.00 | 3.47 | <.001 *** |
|  | Subject | 50 | 421,423 | 8,428.46 | 48.11 | <.001 *** |
|  | Task Day x Subject | 346 | 125,272 | 362.00 | 2.07 | <.001 *** |
|  | Residuals | 1,182 | 207,058 | 175.19 |  |  |
| Energetic | Task Day | 10 | 11,586 | 1,158.60 | 4.81 | <.001 *** |
|  | Subject | 50 | 169,106 | 3,382.12 | 14.03 | <.001 *** |
|  | Task Day x Subject | 346 | 128,011 | 369.95 | 1.54 | <.001 *** |
|  | Residuals | 1,182 | 284,922 | 241.10 |  |  |
| Enjoyment | Task Day | 10 | 2,804 | 280.40 | 1.62 | .097 |
|  | Subject | 50 | 182,900 | 3,658.00 | 21.07 | <.001 *** |
|  | Task Day x Subject | 346 | 108,243 | 312.74 | 1.80 | <.001 *** |
|  | Residuals | 1,182 | 205,232 | 173.68 |  |  |
| Happy | Task Day | 10 | 1,582 | 158.20 | 1.27 | .240 |
|  | Subject | 50 | 139,991 | 2,799.82 | 22.54 | <.001 *** |
|  | Task Day x Subject | 346 | 97,511 | 281.80 | 2.27 | <.001 *** |
|  | Residuals | 1,182 | 146,821 | 124.20 |  |  |
| Motivated | Task Day | 10 | 9,877 | 987.70 | 3.86 | <.001 *** |
|  | Subject | 50 | 161,484 | 3,229.68 | 12.64 | <.001 *** |
|  | Task Day x Subject | 346 | 131,369 | 379.59 | 1.49 | <.001 *** |
|  | Residuals | 1,182 | 302,104 | 255.55 |  |  |
| Relaxed | Task Day | 10 | 6,476 | 647.60 | 3.36 | <.001 *** |
|  | Subject | 50 | 346,832 | 6,936.64 | 35.98 | <.001 *** |
|  | Task Day x Subject | 346 | 135,255 | 390.96 | 2.03 | <.001 *** |
|  | Residuals | 1,182 | 227,889 | 192.81 |  |  |

**Note.** One-way ANOVAs were used to compare self-reported mood scores across participants and task days. $P < 0.10$ ; \* $P < 0.05$ ; \*\* $P < 0.01$ ; \*\*\* $P < 0.001$

### Task-based Affective Bias

Behavioural affective bias was assessed using tasks from the Emotional Test Battery (ETB), a validated measure of mood-related cognitive processing (55, 56), adapted for daily use and optimised for smartphones. Participants completed tasks indexing emotional categorisation (ECAT), emotional recall (EREC) and emotional memory (EMEM) of valenced self-referential words (56). Participants completed one session (which includes ECAT, EREC and EMEM) per evening across six days, with unique word lists on each day. Compared to previous iterations, this version of the EMEM was reduced in length to ensure use of validated word lists and minimise participant burden for daily use. Three affective bias indices were derived: ECAT, EREC and EMEM negative bias. Each index was calculated as the difference between true-negative and true-positive rates (TNR - TPR), such that higher values indicate greater accuracy for negatively valanced stimuli. Participants were included if they completed at least five days of behavioural tasks. Data were excluded when TNR or TPR values for the ECAT and EMEM exceeded 30% or EREC was 0%. A total of 47 participants were included in this analysis. Across each of these measures, descriptive statistics revealed negative or almost-zero values (Fig. 2E-F), with median bias scores of *ECAT negative bias* (median = 0; SD = 8.96), *EREC negative bias* (median = -2.5; SD = 11.42) and *EMEM negative bias* (median = 0; SD = 18.43) being grouped around zero.

As with the self-reported questionnaires, we assessed individual and day-to-day variability in affective bias scores (*ECAT negative bias, EREC negative bias* and *EMEM negative bias*). Analysis showed that individual differences were significant for *ECAT negative bias* (*F*(49, 235)= 3.30, *p*<0.001, η² = 0.41) and *EREC negative bias* (*F*(49)= 1.62, *p*<0.010, η² = 0.237). While the day of assessment was significant for *EREC negative bias* (*F*(5, 235) = 4.00, *p* = 0.002, η² = 0.06) and *EMEM negative bias* (*F*(5, 235) = 4.30, *p*<0.001, η² = 0.069), these effects accounted for considerably less variance overall. Notably, these modest daily fluctuations may reflect variation in the specific words presented on each testing day rather than true changes in affective bias, consistent with the trait-like nature of these measures.

### Relationships Between Task-based Affective Bias and Self-reported Mood Dynamics

To examine how behavioural affective bias relates to self-reported emotion, we tested associations with daily mood ratings and longer-term (weekly) mood characteristics. Behavioural affective bias measures were only weakly associated with same-day mood ratings. Among the bias outcomes, only *ECAT negative bias*, reflecting emotional categorisation bias, showed statistically significant correlations with positive self-reported mood. Greater *ECAT negative bias* was associated with lower ratings of feeling *carefree (R*(45) = -0.14*, p* = 0.02), *energetic* (*R*(45) = -0.13, *p* = 0.03), *enjoyment (R*(45) *=* -0.15*, p =* 0.009), *happy* (*R*(45) = -0.13, *p* = 0.02), and *motivated* (*R*(45) =-0.16, *p* = 0.006). In contrast, bias outcomes that explored emotional recognition and memory showed no reliable relationships with daily mood (*SI Appendix,* Table S3). We next examined whether affective bias was more strongly related to stable weekly mood characteristics. Within this analysis, associations were stronger and more consistent, driven primarily by *ECAT negative bias*. Greater *ECAT negative bias* showed moderate negative correlations with weekly mean levels of *happiness* (*R*(45) = -0.30, *p* = 0.03), *energy* (*R* (45) = -0.37, *p* = 0.009), *motivation* (*R*(45) = -0.36, *p* = 0.01) and *enjoyment* (*R* (45) = -0.31, *p* = 0.03). Again, *EREC negative bias* and *EMEM negative bias* showed minimal or inconsistent relationships with weekly mood outcomes (*SI Appendix,* Table S4).

### Relationships Between the Acute Effects of Light Exposure and Mood Outcomes

After full cleaning for light and mood variables, 48 participants were used in the final analysis. We first examined the short-term impact of light exposure on mood within the 30 min preceding self-reported ratings, based on previous animal research (57, 58). Individual slopes relating melanopic EDI intensity to each mood dimension were predominantly positive, although some participants exhibited negligible effects, possibly indicating variability in light sensitivity (Fig. 3A).

**Fig. 3.**
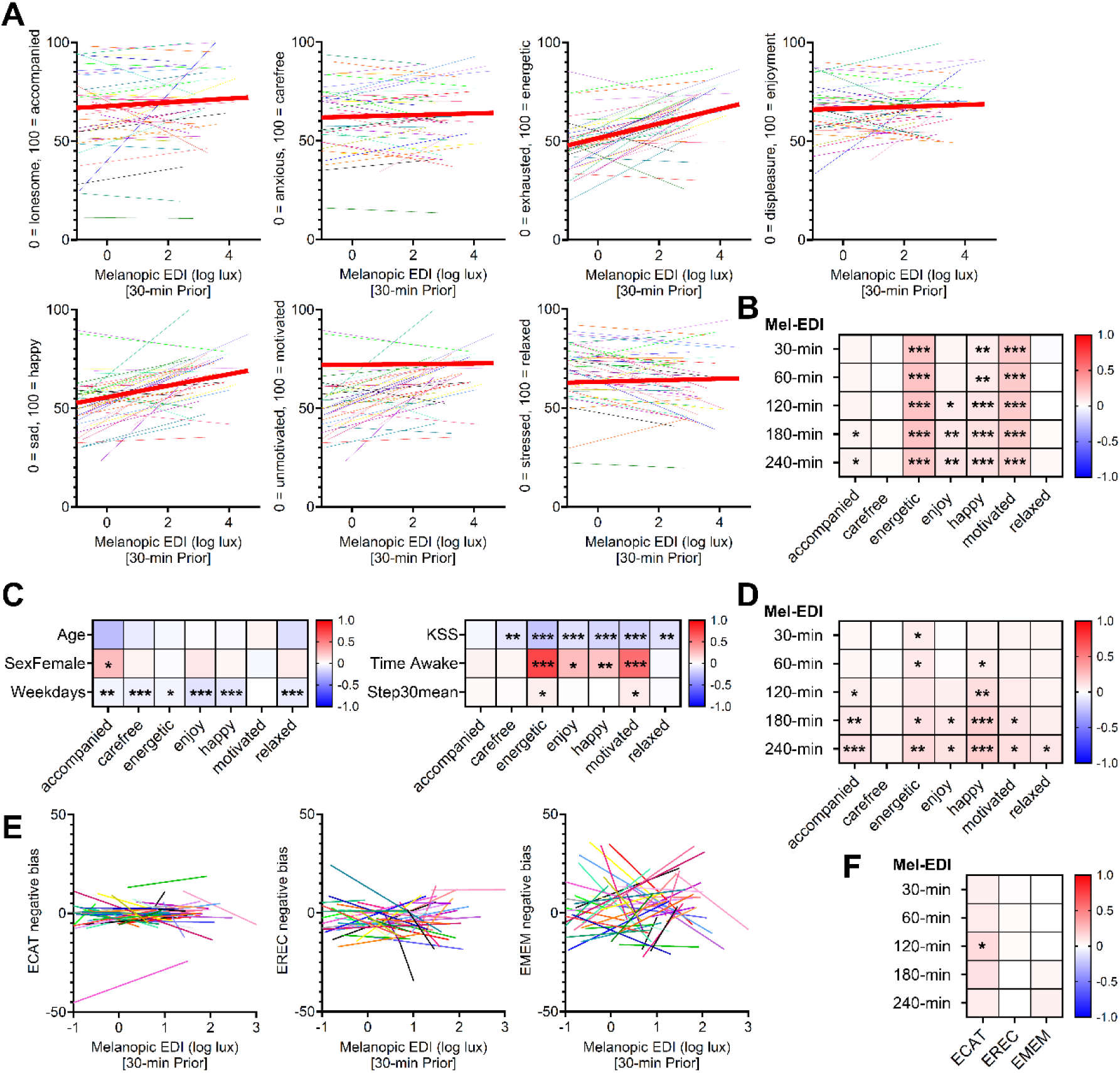
Acute effects of light exposure on mood and affective bias. Relationships between acute light exposure over the preceding 30 min to 4 h and both self-reported mood (*n* = 48) and affective bias (*n* = 47). (A) Individual slopes showing the relationship between mean melanopic equivalent daylight illuminance (melanopic EDI) in the 30 min before assessment and self-reported mood scores. Each line represents one participant, and the mean slope is highlighted in red. (B) Heat map of acute light-exposure effects on mood without adjustment for significant covariates. Models tested mean melanopic EDI in time windows ranging from 30 to 240 min before each mood rating. Standardised effect sizes from linear mixed-effects models are shown, with color indicating direction and magnitude of the association (red, positive; blue, negative). Asterisks indicate significant associations based on Type III ANOVA (P < 0.05, *; P < 0.01, **; P < 0.001, ***). Mood states assessed include accompanied, carefree, energetic, enjoyment, happiness, motivated and relaxed. (C) Heat map of associations between potential covariates and self-reported mood ratings. Covariates include sleep-related measures, such as morning Karolinska Sleepiness Scale (KSS) score, time awake, and steps taken in the preceding 30 min (Step30mean), as well as demographic and contextual variables, including age, sex, and weekday versus weekend. (D) Heat map of acute light-exposure effects on mood after adjustment for significant covariates. Models tested mean melanopic EDI in time windows ranging from 30 to 240 min before each mood rating. Standardised effect sizes from linear mixed-effects models are shown, with color indicating direction and magnitude of the association (red, positive; blue, negative). Asterisks indicate significant associations based on Type III ANOVA (P < 0.05, *; P < 0.01, **; P < 0.001, ***). Mood states assessed include accompanied, carefree, energetic, enjoyment, happiness, motivated, and relaxed. (E) Individual slopes showing the relationship between mean melanopic EDI in the 30 min before completion of the affective bias tasks and affective bias outcomes. Outcomes include ECAT negative bias, EREC negative bias, and EMEM negative bias. Each line represents one participant. (F) Heat map of acute light-exposure effects on affective bias. Standardised effect sizes from linear mixed-effects models are shown, with color indicating direction and magnitude of the association (red, positive; blue, negative), and asterisks indicate significant associations based on Type III ANOVA (P < 0.05, *; P < 0.01, **; P < 0.001, ***). Models tested mean melanopic EDI in time windows ranging from 30 to 240 min before task completion. ECAT, emotional categorisation; EREC, emotional recall; EMEM, emotional memory.

Across the cohort, higher melanopic EDI during the prior 30 min was significantly associated with greater feelings of *energetic* (β = 0.21, 95% CI [0.14, 0.28], *p*<0.001, η² = 0.44), *motivated* (β = 0.20, 95% CI [0.13, 0.27], *p*<0.001, η² = 0.42), and *happy* (β = 0.06, 95% CI [0.01, 0.10], *p* = 0.01, η² = 0.04) moods. Temporal analyses revealed that enjoyment and accompanied mood strengthened with longer exposure windows (up to 4 h), whereas *energetic, motivated* and *happy* moods were most responsive to light in the preceding 30 min (SI Appendix, Table S5, Fig. 3B).

To isolate light-specific effects, models were adjusted for covariates including, morning KSS, time of day, time awake, last 30 minutes mean step count, age, sex, weekday/weekend, photoperiod and chronotype (MCTQ-MSFsc). Morning KSS was negatively associated with *energetic* (*R*(46) = - 0.23, *p* < 0.001) and *motivated* (*R*(46) = - 0.17, *p* < 0.001) moods, whereas time awake positively predicted these energetic (*R*(46) = 0.73, *p* < 0.001) and motivated *(R*(46) = 0.57, *p* < 0.001) states (Fig. 3C). Physical activity (last 30 minutes step mean) showed small but significant positive associations with *energetic* (*R*(48) = 0.06, *p* < 0.001) and *motivated* (*R*(48) = 0.07, *p* < 0.05). (Fig. 3C; SI Appendix, Table S2). Weekends were also linked to generally higher self-reported mood ratings. Notably, after adjustment, the acute light effect, although attenuated, remained detectable, with *energetic* mood still significantly associated with 30-min prior exposure (β = 0.07, 95% CI [0.00, 0.14], *p* = 0.049, η² = 0.07), while associations for other mood dimensions emerged at longer exposure windows (2-4 h; Fig. 3D; SI Appendix, Table S6).

Based on our finding that the preceding 30 minutes was significantly associated with self-reported mood, we applied the same window to examine affective bias. After full cleaning for light and affective bias variables, 47 participants were used in the final analysis. Acute light exposure at 30 minutes showed no significant effects on any affective bias measure, including *ECAT negative bias* (β = 0.06, 95% CI [-0.07, 0.18], *p* = 0.40, η² = <0.001*), EREC negative bias* (β = 0.01, 95% CI [-0.12, 0.14], *p* = 0.87, η² = <0.001) or *EMEM negative bias* (β = -0.005, 95% CI [-0.13, 0.12], *p* = 0.94, η² = <0.001). This relationship continues up to 4 hours, with an anomaly of a significant effect at mean 2 hours of light exposure for *ECAT negative bias* (Fig. 3E-F; Table S7).

### Relationship Between Cumulative Daily Light Exposure and Self-Reported Mood Outcomes

To assess cumulative daily effects, total 24h light exposure above melanopic thresholds of 250 and 1,000 lux melanopic EDI, significantly predicted average mood ratings (n = 48, 7 days). Higher daily time spent above 250 lux melanopic EDI predicted elevated positive self-reported mood scores and related significant for *happiness* (β = 0.15, 95% CI [0.03, 0.26], *F*(1, 34.73) = 6.16, *p* = 0.018, η² = 0.15), *enjoyment* (β = 0.16, 95% CI [0.05, 0.26], F(1, 24.16) = 8.80, *p* = 0.007, η² = 0.27), *accompanied* (β = 0.10, 95% CI [0.01, 0.18], *F*(1, 25.24) = 5.22, *p* = 0.031, η² =0.17) and *relaxed* (β = 0.10, 95% CI [0.00, 0.19], F(1, 291.77) = 4.30, *p* = 0.039, η² = 0.01) moods. Similar relationships were observed for time spent above 1000 lux melanopic EDI (SI Appendix, Table S8, Fig. S4; Fig. 4A). Adjustment for weekday/weekend attenuated time spent above 1000 lux melanopic EDI associations, but time spent above 250 lux melanopic EDI effects remained significant (Fig. 4B; SI Appendix, Table S9).

**Fig. 4.**
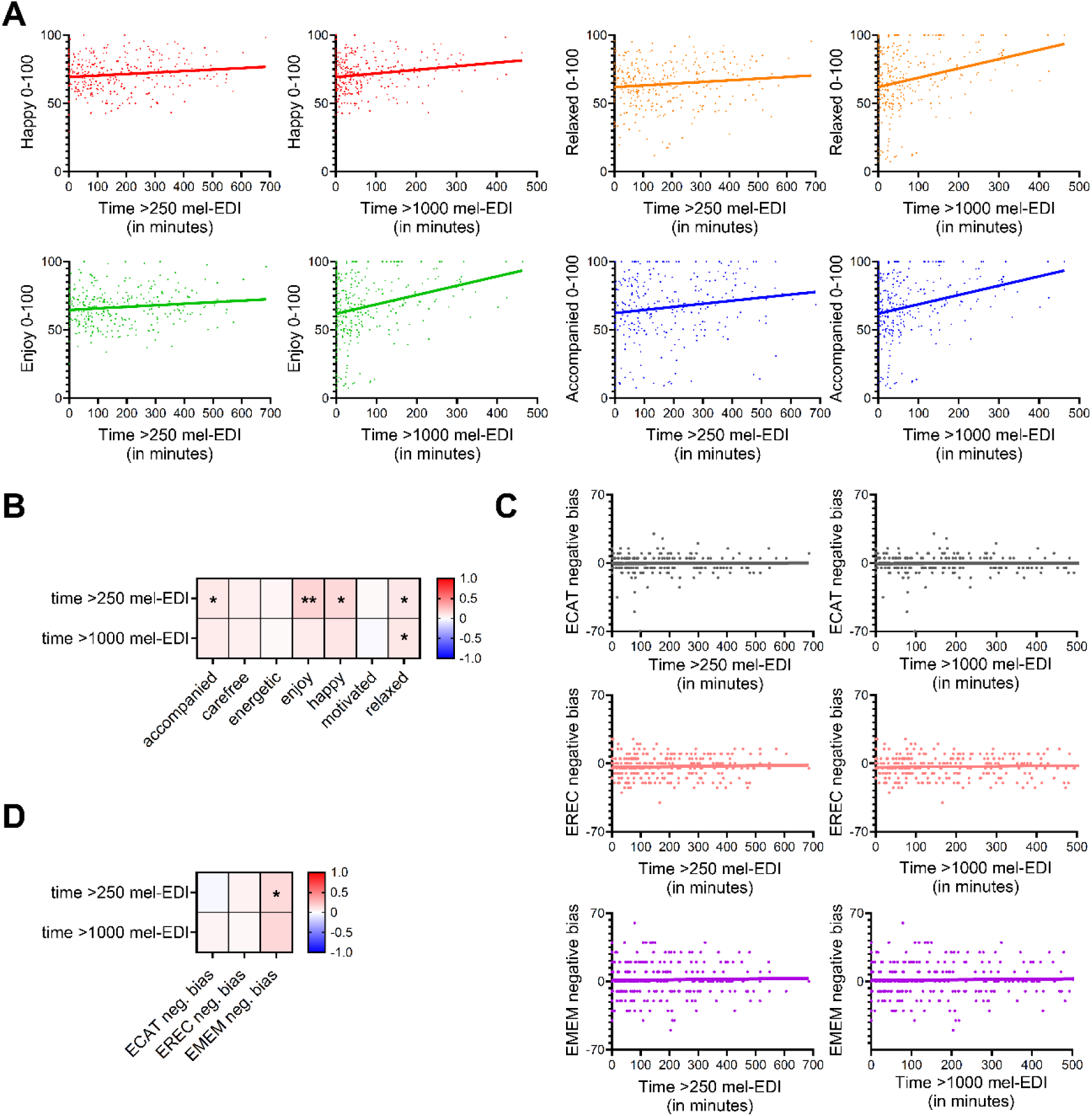
Relationships between daily light exposure and self-reported mood and affective bias. Associations between daily light exposure and self-reported mood (*n* = 48) and affective bias (*n* = 47). Daily light exposure was quantified as time spent above melanopic equivalent daylight illuminance (melanopic EDI) thresholds of 250 lx and 1,000 lx. (**A**) Line graphs showing raw data and mean time spent above 250 lx and 1,000 lx melanopic EDI for mood variables that were significant before covariate adjustment (accompanied, enjoyment, happy, and relaxed). (**B**) Heat map showing associations between daily light exposure and self-reported mood outcomes after adjustment for covariates. Standardised effect sizes indicate the strength and direction of associations (red, positive; white, negative), and asterisks indicate significant associations based on Type III ANOVA (P < 0.05, *; P < 0.01, **; P < 0.001, ***). Daily light exposure variables include time spent above 250 lx melanopic EDI (time >250 mel-EDI) and above 1,000 lx melanopic EDI (time >1000 mel-EDI). Standardised coefficients were derived from linear mixed-effects models, and P values were obtained from Type III ANOVA. (**C**) Line graphs showing raw data and mean time spent above 250 lx and 1,000 lx melanopic EDI for affective bias outcomes, including ECAT negative bias, EREC negative bias, and EMEM negative bias. (**D**) Heat map showing associations between daily light exposure and affective bias outcomes after adjustment for covariates. Standardised effect sizes indicate the strength and direction of associations (red, positive; white, negative), and asterisks indicate significant associations based on Type III ANOVA (P < 0.05, *; P < 0.01, **; P < 0.001, ***). Positive coefficients indicate greater negative bias. ECAT, emotional categorisation; EREC, emotional recall; EMEM, emotional memory.

Analyses of daily light exposure, using the time spent above the thresholds of 250 lx and 1000 lux melanopic EDI metrics (n = 47, 7 days), revealed no significant effects on *ECAT negative bias* (time spent above 250 lux melanopic EDI: β = -0.03, 95% CI [-0.31, 0.25], *F*(1, 21.47) = 0.05, *p* = 0.82, η² = 0.002); time spent above 1000 lux melanopic EDI: β = 0.04, 95% CI [-0.08, 0.17], F(1, 46.31) = 0.51, *p* = 0.48, η² = 0.01) and *EREC negative bias* (time spent above 250 lux melanopic EDI: β = 0.05, 95% CI [-0.09, 0.19], *F*(1, 40.60) = 0.50, *p* = 0.48, η² = 0.01; time spent above 1000 lux melanopic EDI: (β = 0.03, 95% CI [-0.14, 0.20], *F*(1, 1.27) = 0.13, *p* = 0.77, η² = 0.09) (Fig. 4C-D). In contrast, EMEM negative bias showed a significant positive association with time spent above 250 lux melanopic EDI only (β = 0.14, 95% CI [-0.02, 0.27], *p* = 0.02, η² = 0.04), indicating that greater light exposure was associated with increased negative emotional memory bias. These findings suggest that the relationship between bright daily light exposure and emotional memory bias may differ from its generally positive associations with mood and affective state.

## Discussion

Experimental and clinical evidence demonstrates that light influences mood, yet its effects in naturalistic settings remain less well characterised. In particular, it remains unclear which mood dimensions are most sensitive to light and whether these associations reflect acute exposure close to the time of assessment, cumulative exposure across the day or more stable affective processes. Here, we addressed this gap by integrating continuous measures of light exposure, quantified as melanopic EDI, with repeated assessments of self-reported mood and behavioural indices of affective bias in everyday contexts. This approach enabled multimodal sampling of light exposure, physiology and affective state across an ecologically valid timescale.

Our findings revealed that brighter light exposure was consistently associated with more positive self-reported mood. Notably, associations with feelings of *energy, motivation,* and *happiness* emerged following as little as 30 minutes of light exposure preceding assessment, whereas associations with feeling *accompanied* and *enjoyment* were observed after longer durations of exposure (120-180 min). Given that light has well-established alerting effects and that mood is influenced by factors such as physical activity, alertness, chronotype and sleep-related measures, we examined whether these associations persisted after accounting for potential confounding variables. Importantly, several associations remained significant independent of activity levels, alertness and sleep-related measures. Higher melanopic EDI within the preceding 30 minutes was associated with greater feelings of *energy*, suggesting that this relationship is at least partly independent of behavioural arousal. In contrast, associations with other mood dimensions emerged over longer exposure windows. Higher melanopic irradiance was associated with greater feelings of *happiness* after 60 min, *accompaniedness* and *enjoyment* after 120 min, *motivation* after 180 min, and *relaxation* after 240 min.

These findings suggest that different dimensions of mood vary in both their sensitivity to light and the duration of exposure required to influence them. In contrast, greater cumulative daily light exposure above 250 lux melanopic EDI was associated with higher ratings of *happiness*, *enjoyment*, *relaxation*, and feeling *accompanied*, while feelings of *energy* showed no association with cumulative light exposure. Behavioural measures of affective bias, however, showed no robust relationship with light exposure, suggesting that light may be more closely related to state-like mood fluctuations than to more stable cognitive-affective biases in this healthy sample.

Large-scale population studies show that greater daylight exposure is associated with improved mood, sleep and circadian health, as well as reduced risk of depression and anxiety (4, 5). While these studies demonstrate robust links between habitual light environments and mental health, they generally rely on aggregate measures of light exposure and retrospective or clinical assessments of mood. The present study advances this literature by showing that everyday light exposure is associated with within-day variations in mood in healthy individuals. Furthermore, our findings highlight the complexity of the relationship between light exposure and mood, with different affective states showing distinct associations across exposure timescales. These effects are therefore unlikely to be captured by a single total-exposure metric and instead depend on factors such as exposure timing, baseline state, and the temporal window over which light is assessed.

One plausible interpretation is that the acute associations observed here partly reflect light’s established alerting and fatigue-reducing effects. Laboratory studies have shown that bright light can exert acute effects on subjective sleepiness, vigilance and physiological markers of arousal, including waking EEG activity and slow eye movements, with responses varying according to timing, intensity and prior sleep-wake state (46, 59–62). Evidence from experimental studies demonstrates that light can influence a range of non-visual responses, including melatonin secretion, subjective sleepiness, thermoregulation, heart rate, alertness, vigilance and waking EEG activity (29, 63). Applied studies extend this evidence to more naturalistic settings, showing that brighter lighting can improve alertness, vitality, concentration, and aspects of positive mood, while reducing fatigue and irritability, although effects are not uniform across all lighting interventions or outcomes (47, 64, 65–67). These findings are particularly relevant to the present observation that recent light exposure was most consistently associated with feelings of *energy*. However, the associations observed here were not fully explained by measured sleepiness, sleep pressure, or activity levels. Importantly, associations between short-term light exposure and mood persisted after adjusting for morning sleepiness (KSS) and a range of light-related covariates, including time awake, chronotype, recent physical activity, age, sex, and weekday versus weekend effects. These findings suggest that the influence of light on mood cannot be fully attributed to its established effects on alertness or to behavioural factors that typically accompany greater light exposure. Moreover, *energy* and *motivation* showed clear circadian modulation, with both mood dimensions reaching their highest levels around midday, highlighting a strong contribution of time-of-day to positive affect and recent light exposure.

Consistent with this interpretation, human studies indicate that light influences neural and behavioural processes involved in emotion as well as alertness. Light has been shown to enhance task-related responses in brain regions supporting cognition and alertness, while also modulating emotional brain responses and prefrontal-amygdala circuitry in ways associated with reduced negative mood (49, 68–70). The biological mechanisms underlying these associations remain to be determined. However, intrinsically photosensitive retinal ganglion cells (ipRGCs), which express the photopigment melanopsin, are known to contribute to a range of non-visual responses to light, and melanopic EDI provides a biologically informed metric for quantifying light exposure relevant to this pathway (39, 40, 54, 71–73). Animal studies suggest that ipRGC-linked neuronal pathways influence both arousal-related and affective behaviours (57, 58, 74, 75). However, because the present findings cannot be explained solely by subjective alertness, evidence from animal models of mood- and anxiety-related behaviours may be particularly relevant. Nevertheless, such studies provide mechanistic support rather than direct evidence for effects on self-reported mood in humans.

More direct evidence for effects on affect comes from studies showing that morning bright light reduces negative affect in adolescents and that light exposure can decrease the endorsement of negative self-descriptors (76, 77). The present findings may reflect the contribution of multiple interacting mechanisms. Recent light exposure may influence mood through more immediate effects on alertness, arousal, vitality and emotion-related processing, whereas cumulative daily light exposure may act over longer timescales through pathways involving circadian alignment, sleep-wake regulation, repeated exposure to brighter daytime environments and related behavioural contexts. While associations with feelings of *energy* emerged following relatively recent light exposure, other mood dimensions were related to longer periods of exposure, potentially reflecting contributions from both immediate physiological responses and broader influences of daily light history and behavioural context.

Clinical, epidemiological and naturalistic evidence further supports the idea that the effects of light on affective functioning depend on its timing and context rather than total exposure alone. Bright light therapy is an established treatment for seasonal depression and has also shown efficacy in non-seasonal, bipolar, and perinatal depression, demonstrating that appropriately timed light can causally influence mood in clinical populations (10–12, 14, 78, 79). However, the mechanisms underlying these therapeutic effects may differ from those proposed for conventional antidepressant treatments. In a double-blind study of healthy volunteers, a single 60-minute session of morning bright light treatment did not alter affective bias or other measures of emotional information processing assessed immediately afterwards using the Oxford Emotional Test Battery (77). This contrasts with the rapid positive shifts in emotional processing frequently observed following acute antidepressant administration and suggests that the therapeutic effects of bright light may not depend on an immediate modification of affective bias. Instead, such effects may arise through changes in circadian timing, sleep-wake regulation, alertness or other context-dependent processes. This interpretation remains provisional, however, as the study examined the acute effects of a single light exposure in healthy individuals rather than the effects of repeated treatment in a clinical population. More recent naturalistic studies provide a closer parallel to the present findings, showing that environmental bright light, natural light exposure, workday illuminance and morning melanopic light are associated with lower depressive and stress symptoms, reduced momentary exhaustion, improved sleep regularity and higher positive affect, including associations that persist after accounting for physical activity and other behavioural factors (37, 38, 67, 80). Together with the present results, these studies suggest that the relationship between light exposure and mood is shaped by a complex interplay of biological and behavioural factors, with the timing and duration of exposure appearing to be particularly important.

The distinct pattern observed for behavioural measures of affective bias further supports the idea that ambient light may be more closely related to momentary self-reported mood than to more stable cognitive-affective processing in this sample. Affective bias, particularly negative bias on the ECAT, was only weakly associated with momentary self-reported mood but showed stronger relationships with participants’ average weekly mood levels. This suggests that affective bias may primarily reflect more stable, trait-like aspects of emotional processing rather than short-term fluctuations in subjective experience. Consistent with this interpretation, we did not observe robust or consistent associations between light exposure and performance across the emotional test battery, including the ECAT, EREC and EMEM (55, 56). Although some isolated associations emerged, such as a greater negative emotional memory bias on the EMEM task with higher light exposure, these effects were modest and lacked consistency across tasks and exposure windows. This study was conducted in individuals without diagnosed mental health conditions, consistent with baseline assessments indicating generally good mental and physical health. Most participants reported none-to-mild symptoms of depression and anxiety and no participants rated their health as poor. This was further supported by the overall positive bias observed across the emotional test battery. Consequently, the associations observed here may not generalise directly to clinical populations. Individuals with mood disorders, such as depression, may exhibit greater variability in affective processing and altered sensitivity to environmental light exposure, potentially leading to different patterns of light-mood associations. It is also important to note that the ETB was adapted for repeated daily sampling by distributing its components across six consecutive days, rather than administering the full word lists in a single session as originally designed and validated. While necessary for this study, this modification may have altered the psychometric properties of the tasks. Furthermore, reaction time data could not be analysed reliably because of latency variability within the mobile application, limiting analyses to accuracy-based measures. These methodological factors should be considered when interpreting the absence of robust associations between light exposure and affective bias.

Several limitations should be considered. First, the observational design precludes causal inference, and the observed associations may partly reflect reverse causation or unmeasured behavioural and contextual factors that co-vary with light exposure in real-world settings, including outdoor activity, social interaction, occupational schedules and seasonal influences (6, 38). Second, the relatively small, demographically homogeneous sample of healthy young adults may have limited power to detect smaller effects and restricts generalisability to other populations, including adolescents, older adults, shift workers and individuals with mood or sleep disorders. Third, wrist-worn light sensors provide only an indirect estimate of biologically effective ocular light exposure and are susceptible to measurement error arising from device position, clothing coverage and environmental factors. Finally, reliance on self-reported mood measures and the complexity of naturalistic environments make it difficult to fully disentangle the direct effects of light from related influences of sleep, circadian timing, behaviour and social context. Future experimental and semi-naturalistic studies will be important for establishing causality and distinguishing acute from cumulative effects of light exposure on mood.

In conclusion, this study provides evidence that with real-world light exposure, melanopic EDI, is associated with dynamic variations in mood in everyday life. Associations varied across timescales, with recent light exposure relating most strongly to feelings of *energy* and exposure integrated over longer periods relating to a broader range of positive mood states, including *happiness*, *motivation*, *relaxation*, *enjoyment* and feeling *accompanied*. Greater cumulative daily exposure to brighter light was similarly associated with several aspects of positive mood, but not with feelings of *energy*, suggesting that the relationship between light and mood is temporally complex and may involve multiple underlying processes. In contrast, behavioural measures of affective bias showed little evidence of association with light exposure, indicating that everyday light environments may be more closely linked to momentary self-reported mood than to more stable cognitive-affective processing in healthy individuals. Future studies incorporating controlled light manipulations will be important for clarifying the mechanisms underlying these associations, distinguishing rapid from longer-term effects of light and informing the development of targeted light-based strategies to support mental well-being.

## Materials and Methods

### Recruitment

Fifty-one healthy adult volunteers (25 males and 26 females) were recruited into a longitudinal observational study using opportunity sampling. A target sample size of approximately 50 participants was determined based on practical power considerations and recent methodological recommendations for adequately powered perceptual and light-exposure research (81). A total of 51 participants were ultimately recruited, slightly exceeding the planned sample size. Recruitment was conducted through advertisements placed across university buildings, the university website and via word-of-mouth. Participants were recruited across the year, ensuring temporal diversity in light exposure conditions. The study aimed to obtain a healthy but demographically diverse sample to reflect the general population. Eligibility criteria included: (1) aged 18 years or older and residing in the UK, (2) native English speaker, (3) not engaged in night shift work, (4) no recent travel across time zones (within two weeks of participation), (5) ownership of a functioning smartphone without screen damage (to ensure accurate survey responses), and (6) no current diagnosis of depression based on participant knowledge. In addition to the light and mood protocol, participants contributed data on sleep, physical and mental health, which may impose limitations on the specificity of mood-related associations. Participants received financial compensation upon successful completion of the study protocol. The study ran from November 2023 to September 2024 and received ethical approval from the University of Manchester Research Ethics Committee (Reference: 2023-18265-31575).

All participants provided written informed consent at the baseline visit and were reminded of their right to withdraw at any time during the seven-day study period. Participants were instructed to complete the following: (1) a battery of baseline questionnaires, (2) wear a Condor Instruments ActLumus light monitor and Fitbit Charge 5 device, (3) complete five mood questionnaires daily, and (4) complete a morning sleep diary.

### Devices

Participants were provided with an ActLumus light-sensing device (34), which continuously recorded environmental light exposure at one-minute intervals. The device captured data on melanopic EDI. Melanopic EDI is especially important, as it reflects light effects on non-image-forming visual pathways implicated in circadian and mood regulation. The ActLumus sensor has a measurement range of 1–100,000 lux and a 10% accuracy at 1,000 lux, achieved through the integration of ten light channels. Condor devices were worn only during waking hours to prevent obstruction during sleep and were not issued with chargers, as the battery lifespan exceeded the study duration. Participants were advised to avoid wearing long sleeves or garments that could obstruct sensor function. Participants were also provided with a Fitbit Charge 5 wristband sleep and activity tracker (Fitbit, US) (82), to wear continuously each day, including during the night. This collected continuous heart rate, step count, sleep, oxygen saturation and skin temperature. The devices were preconfigured by the researcher and deactivated upon return. Participants were instructed not to alter settings and to wear the device as part of their daily routine.

### Self-reported mood questionnaires

All surveys were administered via Qualtrics (Qualtrics, Provo, UT) (83) and optimised for mobile completion. Baseline assessments captured demographic data (e.g., age, gender, ethnicity), lifestyle behaviours (e.g., smoking, alcohol use), and any relevant health conditions, including psychological, ocular, sleep, or neurological diagnoses. Participants completed validated instruments measuring mood, sleep, and chronotype, including the Generalised Anxiety Disorder-7 (GAD-7) (84), Patient Health Questionnaire-9 (PHQ-9) (85), PROMIS Anxiety short form (NIH), PROMIS Depression short form (NIH), PROMIS Sleep Disturbance short form (NIH), PROMIS Sleep Impairment short form (NIH), and PROMIS Positive Affect and Well-being scales (NIH) (86-90). Chronotype was assessed via the Munich Chronotype Questionnaire (MCTQ) (91).

Daily sleep diaries (using the KSS) were completed each morning to assess self-reported sleep-wake timing, in addition to several times during the day to measure alertness. Mood was assessed five times per day using a digital adaptation of the Visual Analogue Mood Scale (VAMS) (92). Participants rated seven affective states (*Accompanied, Carefree, Energetic, Enjoyment, Happy, Motivated* and *Relaxed*) on a 0–100 visual analogue scale using a slider interface.

### Affective bias tasks

Tasks were also administered via Qualtrics and optimised for mobile completion. Three components of the Emotional Test Battery (EBT) - the Emotional Categorisation Task (ECAT), Emotional Recall Task (EREC) and Emotional Memory Task (EMEM) - were conducted in the evenings (around 20:00) to assess affective state. A practice of these tasks was also completed during the baseline study, giving the participants a chance to ask questions. In contrast to typical administration for a single visit, or visits 7 days or more apart, here tasks were adapted to measure emotional bias across 6 days to reflect affective variation over the study period. Recall of positive vs negative words was measured and correct word recall. Full task instructions are given at the start of the task. The ECAT focuses on emotional bias in response to emotional stimuli. It uses valenced words, based on personality characteristics, e.g., happy, and participants assess whether they would like or dislike to be associated with that word. Participants are instructed to answer as quickly and accurately as they can. Participants were presented with 40 words daily: 50% positive and 50% negative. Each word is presented once across all six sessions. These words are matched in terms of word length and ratings of frequency and imageability. There is an intertrial-interval (response period) which follows stimulus offset during which the participant is required to press a key to identify whether that personality characteristic is one they would “like” or “dislike” to be described as. Participants are marked based on how many of the words they categorise correctly. After this task, participants undertake 2 distraction tasks: a 30 second wait and 4 simple numerical equations, one of each function, e.g. 4x6. This is designed to ensure the participants forget some of the words from the ECAT. The EREC is designed to follow the ECAT, looking at free recall of emotive stimuli presented to the participant. The individual is given as much time as is needed for them to recall words and type these into the provided textbox, which is somewhat different to its typical use of having 3-5 minute timers as a limit. The accuracy of these words is compared to the list presented. These words are assessed on whether they are biased to positive or negative words.

Lastly, the EMEM is like the ECAT in presentation, yet the EMEM assesses recognition, presenting familiar words (from the ECAT) and novel words. The participant must decide if the words were present or absent in the ECAT. Words are randomly presented, and novel words are matched with the familiar words for word length and their meaning. In comparison to typical EMEM iterations which include 80 words (40 familiar, 40 novel – each 50% positive and 50% negative), we limited to 20 words (10 familiar, 10 novel - each 50% positive and 50% negative) to enable use of validated word lists across six days and to minimise task length for daily completion. For this task, overall affective accuracy bias scores are measured, measuring their negative accuracy – positive accuracy to get a bias score.

### Study Procedure

Following initial contact, participants attended a baseline session at the University of Manchester. During this session, the study protocol was explained in detail and written informed consent was obtained. Participants were issued the ActLumus device and received detailed instructions on usage. The Qualtrics platform was introduced and participants were provided with a reference guide and contact information for troubleshooting. Participants then completed the full set of baseline surveys and affective bias tasks as described above.

From Day 2 to Day 7, participants were instructed to wear the light monitor device during all waking hours, complete a sleep diary each morning and complete five mood surveys per day at distributed time points. Participants were encouraged to complete as many self-reported mood entries as possible. They were also asked to complete behavioural affective bias tasks in the evening (around 20:00), when they had little distraction and could dedicate 20 minutes of their attention to it. A visual schedule (SI Appendix, Table. S10) was provided to facilitate compliance.

On Day 8, participants were instructed to complete one final sleep diary and two mood assessments before returning the devices. If a full day of data was missed during the study period, participants were asked to extend their participation to achieve a full 7-day dataset.

### Statistical analysis

All data cleaning, statistical analyses and plotting were conducted using RStudio (R version 4.4.1) (93) and GraphPad Prism (Version 10.2.2) (94). For all descriptive and inferential analyses, log10-transformed melanopic EDI values were used. Descriptive statistics were reported as means and standard deviations for continuous variables, and as frequencies with percentages for categorical variables. For exploratory comparisons, mean differences in mood scores and light exposure variables were assessed using independent t-tests or Pearson’s coefficients.

To assess the variance in self-reported mood scores explained by task day and individual participants, a one-way ANOVA with an interaction term was conducted. Variance explained in behavioural mood task scores by task day and individual participants was also assessed using a one-way ANOVA. To investigate within-subject variation in self-reported mood scores, we computed the median and standard deviation of scores per participant and fitted a cosine model with a 24-hour period to adjust for diurnal patterns, extracting metrics such as mean absolute residuals, mesor, and amplitude. Median, standard deviation, mesor, amplitude, and residuals were compared between self-reported mood categories using one-way ANOVA followed by Tukey post-hoc tests. To assess long-term correlation between self-reported mood and affective bias tasks, Weekly mood patterns were modelled using cosine functions to derive two rhythm parameters: the mesor, representing each participant’s average weekly mood level and the amplitude, reflecting the extent of weekly mood variation. Bias scores were averaged within participants to match this longer timescale.

Several covariates were included to assess their associations with mood outcomes: morning KSS (subjective sleepiness measured using the Karolinska Sleepiness Scale within two hours of waking), clock time of day (with the day defined as starting at 04:00), time awake in hours, mean step count in the 30 minutes preceding mood reporting, weekday versus weekend, chronotype (MCTQ-MSFsc), photoperiod duration, and participant age and sex. Each covariate was individually tested for association with self-reported mood scores using linear mixed-effects models including random intercepts and slopes for participants. Time of day was modelled using harmonic (sine and cosine) terms, and time awake was modelled using quadratic terms. These models were implemented using the lme4 (v 2.0-1) and lmerTest (v 3.2-1) packages in R. In addition, the association between time of day and light exposure was evaluated using a nonlinear cosine regression model. All fixed-effect coefficients in mixed models were standardised by scaling predictors and outcomes by their sample standard deviations. This rescaling enabled comparison of standardised coefficients across different mood outcomes, which were reported with 95% confidence intervals. Eta-squared (η²) and R^2^ (conditional and marginal) were calculated as a measure of effect size. Standardisation and effect size parameters were calculated using performance (v0.17.0), parameters (v0.29.1), and effectsize (v1.0.2) packages in R.

Light exposure prior to self-reported or behavioural mood assessments was examined using two approaches. First, recent light exposure was quantified as the average lux melanopic EDI in the 30-minute window preceding each mood report. Second, light exposure history was defined as the average lux melanopic EDI at 60, 120, 180, and 240 minutes prior to mood assessments. Daily light exposure was characterised by calculating the total time spent above melanopic thresholds of 250 lux melanopic EDI and 1000 lux melanopic EDI, measured in minutes when full daily exposure measurements were available (<30 min gaps per day). Associations between recent light exposure (mean light level in the previous 30 minutes) and self-reported mood scores were examined using linear mixed-effects models with participant-level random intercepts and slopes. Fully adjusted models were then constructed by including covariates that showed significant associations with self-reported mood scores, in order to account for potential confounding effects of time of day, time awake, morning KSS, mean step count in the previous 30 minutes, and weekday versus weekend. Similarly, associations between recent light exposure and behavioural mood scores were examined using unadjusted linear mixed-effects models with participant-level random intercepts and slopes. Historical light exposure variables (up to four hours prior to mood reporting) were analysed using both unadjusted and fully adjusted linear mixed-effects models for self-reported mood scores. Associations between light history variables and behavioural mood scores were also examined using unadjusted models. Finally, daily average mood scores per participant were computed and their associations with daily light exposure metrics (time spent above 250 and 1000 lux melanopic EDI) were examined using linear mixed-effects models, followed by additional models adjusted for weekday versus weekend. Finally, daily light exposure metrics were compared with behavioural affective bias scores.

## Data, Materials, and Software Availability

Software and hardware designs of the wearable light dosimeter data have been deposited in GitHub (XXX (95).

## Acknowledgments

This work was supported by the Biotechnology and Biological Sciences Research Council (BBSRC) Collaborative Awards in Science and Engineering (CASE) PhD Studentship with Signify to N.M., B.B.O. and C.R. A.L.G and C.J.H are supported through the National Institute for Health and Care Research (NIHR) Oxford Health Biomedical Research Centre (BRC). The views expressed are those of the author(s) and not necessarily those of the NIHR or the Department of Health and Social Care.

## Author contributions

C.R., A.D., A.L.G., C.J.H., B.B.O., and N.M. Designed research: C.R., A.D., B.B.O., and N.M., provided methodology: A.L.G., and C.J.H., performed research: C.R., contributed new analytic tools: A.D., analysed data: A.D., and A.E. visualised the data: C.R., wrote the paper: C.R., A.D., and N.M.

## References

1. C. Cajochen, et al., Evidence That Homeostatic Sleep Regulation Depends on Ambient Lighting Conditions during Wakefulness. Clocks Sleep 1, 517–531 (2019).

2. C. Blume, C. Garbazza, M. Spitschan, Effects of light on human circadian rhythms, sleep and mood. Somnologie (Berl) 23, 147–156 (2019).

3. D. Stewart, U. Albrecht, Beyond vision: effects of light on the circadian clock and mood-related behaviours. NPJ Biol Timing Sleep 2, 12 (2025).

4. A. C. Burns, et al., Time spent in outdoor light is associated with mood, sleep, and circadian rhythm-related outcomes: A cross-sectional and longitudinal study in over 400,000 UK Biobank participants. J Affect Disord 295, 347–352 (2021).

5. A. C. Burns, et al., Day and night light exposure are associated with psychiatric disorders: an objective light study in >85,000 people. Nat. Mental Health 1, 853–862 (2023).

6. A. Didikoglu, et al., Associations between light exposure and sleep timing and sleepiness while awake in a sample of UK adults in everyday life. Proc Natl Acad Sci U S A 120, e2301608120 (2023).

7. L. Schlangen, et al., Situation analysis on lighting for health and well-being. [Preprint] (2014). Available at: https://www.researchgate.net/doi/10.13140/RG.2.1.5117.0166 [Accessed 24 June 2026].

8. B. Bano-Otalora, et al., Bright daytime light enhances circadian amplitude in a diurnal mammal. Proc. Natl. Acad. Sci. U.S.A. 118, e2100094118 (2021).

9. Global, regional, and national burden of 12 mental disorders in 204 countries and territories, 1990–2019: a systematic analysis for the Global Burden of Disease Study 2019. The Lancet Psychiatry 9, 137–150 (2022).

10. A. Wirz-Justice, et al., Chronotherapeutics (light and wake therapy) in affective disorders. Psychol Med 35, 939–944 (2005).

11. D. A. Oren, et al., An open trial of morning light therapy for treatment of antepartum depression. Am J Psychiatry 159, 666–669 (2002).

12. C. Garbazza, et al., Sustained remission from perinatal depression after bright light therapy: A pilot randomised, placebo-controlled trial. Acta Psychiatr Scand 146, 350–356 (2022).

13. N. E. Rosenthal, et al., Seasonal affective disorder. A description of the syndrome and preliminary findings with light therapy. Arch Gen Psychiatry 41, 72–80 (1984).

14. R. W. Lam, et al., Efficacy of Bright Light Treatment, Fluoxetine, and the Combination in Patients With Nonseasonal Major Depressive Disorder: A Randomized Clinical Trial. JAMA Psychiatry 73, 56 (2016).

15. M. Spitschan, Time-Varying Light Exposure in Chronobiology and Sleep Research Experiments. Front. Neurol. 12, 654158 (2021).

16. C. Blume, C. Garbazza, M. Spitschan, Effects of light on human circadian rhythms, sleep and mood. Somnologie (Berl) 23, 147–156 (2019).

17. L. J. M. Schlangen, L. L. A. Price, The Lighting Environment, Its Metrology, and Non-visual Responses. Front. Neurol. 12, 624861 (2021).

18. C. A. Czeisler, et al., Bright Light Induction of Strong (Type 0) Resetting of the Human Circadian Pacemaker. Science 244, 1328–1333 (1989).

19. D. B. Boivin, J. F. Duffy, R. E. Kronauer, C. A. Czeisler, Dose-response relationships for resetting of human circadian clock by light. Nature 379, 540–542 (1996).

20. J. M. Zeitzer, et al., Temporal dynamics of late-night photic stimulation of the human circadian timing system. American Journal of Physiology-Regulatory, Integrative and Comparative Physiology 289, R839–R844 (2005).

21. J. F. Duffy, C. A. Czeisler, Effect of Light on Human Circadian Physiology. Sleep Medicine Clinics 4, 165–177 (2009).

22. S. B. S. Khalsa, M. E. Jewett, C. Cajochen, C. A. Czeisler, A Phase Response Curve to Single Bright Light Pulses in Human Subjects. The Journal of Physiology 549, 945–952 (2003).

23. D. W. Rimmer, et al., Dynamic resetting of the human circadian pacemaker by intermittent bright light. American Journal of Physiology-Regulatory, Integrative and Comparative Physiology 279, R1574–R1579 (2000).

24. C. Gronfier, K. P. Wright, R. E. Kronauer, M. E. Jewett, C. A. Czeisler, Efficacy of a single sequence of intermittent bright light pulses for delaying circadian phase in humans. American Journal of Physiology-Endocrinology and Metabolism 287, E174–E181 (2004).

25. A. Chang, et al., Human responses to bright light of different durations. The Journal of Physiology 590, 3103–3112 (2012).

26. S. A. Rahman, et al., Circadian phase resetting by a single short-duration light exposure. JCI Insight 2, e89494 (2017).

27. G. C. Brainard, et al., Action Spectrum for Melatonin Regulation in Humans: Evidence for a Novel Circadian Photoreceptor. J. Neurosci. 21, 6405–6412 (2001).

28. K. Thapan, J. Arendt, D. J. Skene, An action spectrum for melatonin suppression: evidence for a novel non-rod, non-cone photoreceptor system in humans. The Journal of Physiology 535, 261–267 (2001).

29. S. W. Lockley, G. C. Brainard, C. A. Czeisler, High Sensitivity of the Human Circadian Melatonin Rhythm to Resetting by Short Wavelength Light. The Journal of Clinical Endocrinology & Metabolism 88, 4502–4505 (2003).

30. S. L. Chellappa, et al., Acute exposure to evening blue-enriched light impacts on human sleep. Journal of Sleep Research 22, 573–580 (2013).

31. C. Cajochen, et al., High Sensitivity of Human Melatonin, Alertness, Thermoregulation, and Heart Rate to Short Wavelength Light. The Journal of Clinical Endocrinology & Metabolism 90, 1311–1316 (2005).

32. M. Münch, et al., Wavelength-dependent effects of evening light exposure on sleep architecture and sleep EEG power density in men. American Journal of Physiology-Regulatory, Integrative and Comparative Physiology 290, R1421–R1428 (2006).

33. N. Mohammadian, et al., A Wrist-Worn Internet of Things Sensor Node for Wearable Equivalent Daylight Illuminance Monitoring. IEEE Internet Things J. 11, 16148–16157 (2024).

34. ActTrust wrist-worn actigraphy device. (2025). Deposited 2025.

35. M. Gardesevic, et al., Brighter Time: A Smartphone App Recording Cognitive Task Performance and Illuminance in Everyday Life. Clocks Sleep 4, 577–594 (2022).

36. A. Didikoglu, et al., Relationships between light exposure and aspects of cognitive function in everyday life. Commun Psychol 4, 5 (2025).

37. D. A. Wallace, S. Redline, T. Sofer, J. Kossowsky, Environmental Bright Light Exposure, Depression Symptoms, and Sleep Regularity. JAMA Netw Open 7, e2422810 (2024).

38. R. J. Fitton, et al., Naturalistic light exposure patterns in relation to medication status, mood symptoms, and chronotype. npj Biol Timing Sleep 3, 5 (2026).

39. R. J. Lucas, et al., Measuring and using light in the melanopsin age. Trends in Neurosciences 37, 1–9 (2014).

40. I. Provencio, G. Jiang, W. J. De Grip, W. P. Hayes, M. D. Rollag, Melanopsin: An opsin in melanophores, brain, and eye. Proc Natl Acad Sci U S A 95, 340–345 (1998).

41. D. B. Boivin, Complex Interaction of the Sleep-Wake Cycle and Circadian Phase Modulates Mood in Healthy Subjects. Arch Gen Psychiatry 54, 145 (1997).

42. T. A. LeGates, D. C. Fernandez, S. Hattar, Light as a central modulator of circadian rhythms, sleep and affect. Nat Rev Neurosci 15, 443–454 (2014).

43. N. Milosavljevic, How Does Light Regulate Mood and Behavioral State? Clocks Sleep 1, 319–331 (2019).

44. C. Baglioni, et al., Insomnia as a predictor of depression: A meta-analytic evaluation of longitudinal epidemiological studies. Journal of Affective Disorders 135, 10–19 (2011).

45. P. K. Alvaro, R. M. Roberts, J. K. Harris, A Systematic Review Assessing Bidirectionality between Sleep Disturbances, Anxiety, and Depression. Sleep 36, 1059–1068 (2013).

46. P. Badia, B. Myers, M. Boecker, J. Culpepper, J. R. Harsh, Bright light effects on body temperature, alertness, EEG and behavior. Physiology & Behavior 50, 583–588 (1991).

47. K. C. H. J. Smolders, Y. A. W. De Kort, Bright light and mental fatigue: Effects on alertness, vitality, performance and physiological arousal. Journal of Environmental Psychology 39, 77–91 (2014).

48. C. Cajochen, Alerting effects of light. Sleep Medicine Reviews 11, 453–464 (2007).

49. G. Vandewalle, et al., Spectral quality of light modulates emotional brain responses in humans. Proc Natl Acad Sci U S A 107, 19549–19554 (2010).

50. D. C. Fernandez, et al., Light Affects Mood and Learning through Distinct Retina-Brain Pathways. Cell 175, 71–84.e18 (2018).

51. C. J. Harmer, et al., Effect of Acute Antidepressant Administration on Negative Affective Bias in Depressed Patients. AJP 166, 1178–1184 (2009).

52. B. R. Godlewska, M. Browning, R. Norbury, P. J. Cowen, C. J. Harmer, Early changes in emotional processing as a marker of clinical response to SSRI treatment in depression. Transl Psychiatry 6, e957–e957 (2016).

53. N. T. M. Huneke, et al., Placebo effects in randomized trials of pharmacological and neurostimulation interventions for mental disorders: An umbrella review. Mol Psychiatry 29, 3915–3925 (2024).

54. M. Spitschan, et al., Verification, analytical validation and clinical validation (V3) of wearable dosimeters and light loggers. Digit Health 8, 20552076221144858 (2022).

55. F. F. Ramli, et al., Negative bias in encoding and recall memory in depressed patients with inadequate response to antidepressant medication. Psychopharmacology (Berl) 243, 513–520 (2026).

56. C. J. Harmer, et al., Agomelatine facilitates positive versus negative affective processing in healthy volunteer models. J Psychopharmacol 25, 1159–1167 (2011).

57. N. Milosavljevic, J. Cehajic-Kapetanovic, C. A. Procyk, R. J. Lucas, Chemogenetic Activation of Melanopsin Retinal Ganglion Cells Induces Signatures of Arousal and/or Anxiety in Mice. Curr Biol 26, 2358–2363 (2016).

58. G. Wang, et al., Short-term acute bright light exposure induces a prolonged anxiogenic effect in mice via a retinal ipRGC-CeA circuit. Sci Adv 9, eadf4651 (2023).

59. C. Cajochen, Alerting effects of light. Sleep Medicine Reviews 11, 453–464 (2007).

60. C. Cajochen, J. M. Zeitzer, C. A. Czeisler, D.-J. Dijk, Dose-response relationship for light intensity and ocular and electroencephalographic correlates of human alertness. Behavioural Brain Research 115, 75–83 (2000).

61. J. Phipps-Nelson, J. R. Redman, D.-J. Dijk, S. M. W. Rajaratnam, Daytime Exposure to Bright Light, as Compared to Dim Light, Decreases Sleepiness and Improves Psychomotor Vigilance Performance. Sleep 26, 695–700 (2003).

62. J. L. Souman, A. M. Tinga, S. F. Te Pas, R. Van Ee, B. N. S. Vlaskamp, Acute alerting effects of light: A systematic literature review. Behavioural Brain Research 337, 228–239 (2018).

63. C. Cajochen, et al., High Sensitivity of Human Melatonin, Alertness, Thermoregulation, and Heart Rate to Short Wavelength Light. The Journal of Clinical Endocrinology & Metabolism 90, 1311–1316 (2005).

64. A. U. Viola, L. M. James, L. J. Schlangen, D.-J. Dijk, Blue-enriched white light in the workplace improves self-reported alertness, performance and sleep quality. Scand J Work Environ Health 34, 297–306 (2008).

65. M. G. Figueiro, L. Sahin, B. Wood, B. Plitnick, Light at Night and Measures of Alertness and Performance: Implications for Shift Workers. Biological Research For Nursing 18, 90–100 (2016).

66. A. Borisuit, F. Linhart, J.-L. Scartezzini, M. Münch, Effects of realistic office daylighting and electric lighting conditions on visual comfort, alertness and mood. Lighting Research & Technology 47, 192–209 (2015).

67. S. Frick, L. van der Meij, K. Smolders, E. Demerouti, Y. de Kort, The effect of time and day of the week on burnout-related experiences: an experience sampling study. European Journal of Work and Organizational Psychology 33, 276–293 (2024).

68. G. Vandewalle, et al., Daytime light exposure dynamically enhances brain responses. Curr Biol 16, 1616–1621 (2006).

69. A. Alkozei, et al., Exposure to Blue Wavelength Light Is Associated With Increases in Bidirectional Amygdala-DLPFC Connectivity at Rest. Front Neurol 12, 625443 (2021).

70. A. Alkozei, et al., Exposure to Blue Light Increases Subsequent Functional Activation of the Prefrontal Cortex During Performance of a Working Memory Task. Sleep 39, 1671–1680 (2016).

71. H. Bailes, R. Lucas, Human melanopsin forms a pigment maximally sensitive to blue light (max 479 nm) supporting activation of Gq/11 and Gi/o signalling cascades. Proceedings. Biological sciences / The Royal Society 280, 20122987 (2013).

72. T. M. Brown, et al., Recommendations for daytime, evening, and nighttime indoor light exposure to best support physiology, sleep, and wakefulness in healthy adults. PLoS Biol 20, e3001571 (2022).

73. S. Hattar, H.-W. Liao, M. Takao, D. M. Berson, K.-W. Yau, Melanopsin-Containing Retinal Ganglion Cells: Architecture, Projections, and Intrinsic Photosensitivity. Science 295, 1065–1070 (2002).

74. L. Lazzerini Ospri, G. Prusky, S. Hattar, Mood, the Circadian System, and Melanopsin Retinal Ganglion Cells. Annu. Rev. Neurosci. 40, 539–556 (2017).

75. D. C. Fernandez, et al., Light Affects Mood and Learning through Distinct Retina-Brain Pathways. Cell 175, 71–84.e18 (2018).

76. J. Kopřivová, et al., Bright light exposure reduces negative affect and modulates EEG activity in sleep-deprived and well-rested adolescents. Front. Behav. Neurosci. 19 (2025).

77. M. T. Burge, et al., Blue light influences negative thoughts of self. SLEEP 48, zsaf034 (2025).

78. R. N. Golden, et al., The Efficacy of Light Therapy in the Treatment of Mood Disorders: A Review and Meta-Analysis of the Evidence. AJP 162, 656–662 (2005).

79. D. K. Sit, et al., Adjunctive Bright Light Therapy for Bipolar Depression: A Randomized Double-Blind Placebo-Controlled Trial. AJP 175, 131–139 (2018).

80. S. A. Shankman, et al., Disentangling the effects of daily physical activity and natural white light exposure on affect. Journal of Psychopathology and Clinical Science 134, 520–526 (2025).

81. J. Zauner, L. Udovicic, M. Spitschan, Power analysis for personal light exposure measurements and interventions. PLoS ONE 19, e0308768 (2024).

82. Fitbit Charge Fitness Tracker. (2023). Deposited 2023.

83. Qualtrics. (2026). Deposited 2026.

84. R. L. Spitzer, K. Kroenke, J. B. W. Williams, B. Löwe, A brief measure for assessing generalized anxiety disorder: the GAD-7. Arch Intern Med 166, 1092–1097 (2006).

85. K. Kroenke, R. L. Spitzer, J. B. W. Williams, The PHQ-9: Validity of a brief depression severity measure. J Gen Intern Med 16, 606–613 (2001).

86. HealthMeasures (2025) PROMIS anxiety short form 8a v1.0. National Institutes of Health.

87. HealthMeasures (2025) PROMIS depression short form 8a v1.0. National Institutes of Health.

88. HealthMeasures (2025) PROMIS positive affect short form v1.0. National Institutes of Health.

89. HealthMeasures (2025) PROMIS sleep disturbance short form 8b v1.0. National Institutes of Health.

90. HealthMeasures (2025) PROMIS sleep-related impairment short form 8a v1.0. National Institutes of Health.

91. T. Roenneberg, A. Wirz-Justice, M. Merrow, Life between Clocks: Daily Temporal Patterns of Human Chronotypes. J Biol Rhythms 18, 80–90 (2003).

92. G. D. van Rijsbergen, C. L. H. Bockting, M. Berking, M. W. J. Koeter, A. H. Schene, Can a one-item mood scale do the trick? Predicting relapse over 5.5-years in recurrent depression. PLoS One 7, e46796 (2012)

93. R Core Team (2024) R: A language and environment for statistical computing (version 4.4.1). R Foundation for Statistical Computing, Vienna.

94. GraphPad Software (2024) GraphPad Prism. GraphPad Software, San Diego.

95. Roddis C, Didikoglu A, Ebrahimi A, Gillespie A, Harmer C, Bano Otalora B, Milosavljevic N (2026) Analysis scripts for differential effects of acute and daily light exposure on positive mood in a non-depressed cohort. GitHub repository.

