## Supplementary information for "Acute and cumulative light exposure differentially influence mood in healthy adults"

**This PDF file includes:**

Figures S1 to S4
Tables S1 to S10

**Other supporting materials for this manuscript include the following:**

Software S1 to Sx

**Figures**

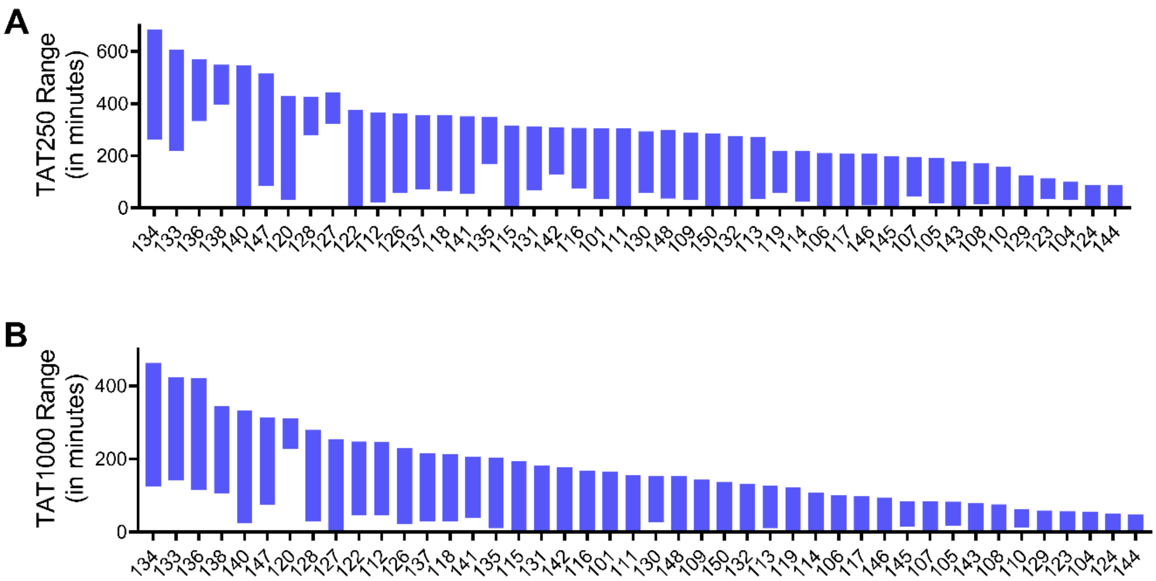


**Fig. S1. Daily average time above 250 and 1,000 lux melanopic equivalent daylight illuminance (EDI) across participants (n=49).** The figure shows the mean daily duration, in minutes, that each participant was exposed to melanopic EDI levels exceeding (A) 250 lux (TAT250) and (B) 1,000 lux (TAT1000). Participants are ordered in descending order of exposure duration within each panel.

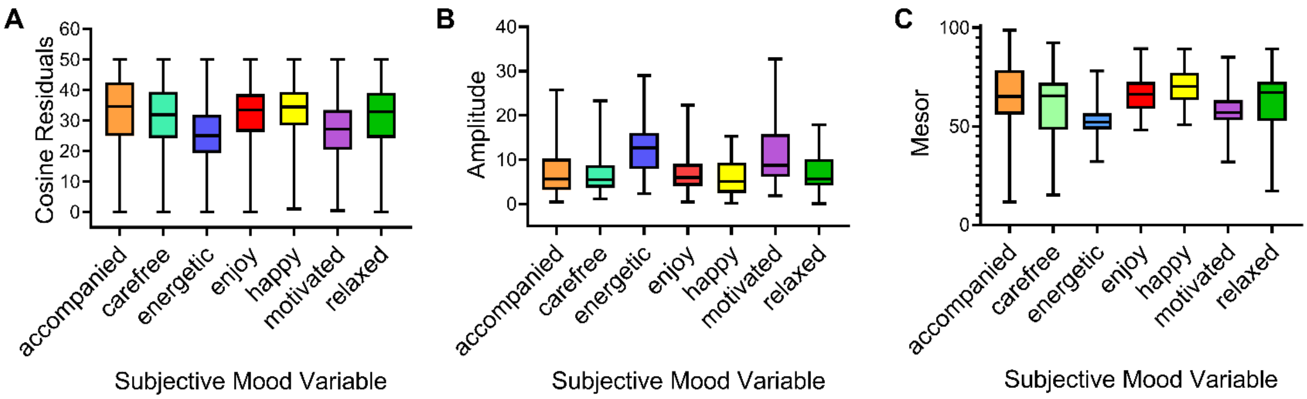


**Fig. S2. Variation in self-reported mood scores and daily rhythmicity (n=50).** Data from all participants were analysed by calculating participant-level summary statistics for each of seven mood states-feeling *accompanied, carefree, energetic, enjoyment, happiness, motivation,* and *relaxation*, and fitting cosine models to assess time-of-day variation. (A) Residual error from the fitted cosine models, with lower values indicating a better fit and stronger rhythmicity. (B) Amplitude of the fitted rhythm, representing the magnitude of daily mood variation. (C) Mesor, representing the mean mood level across the day.

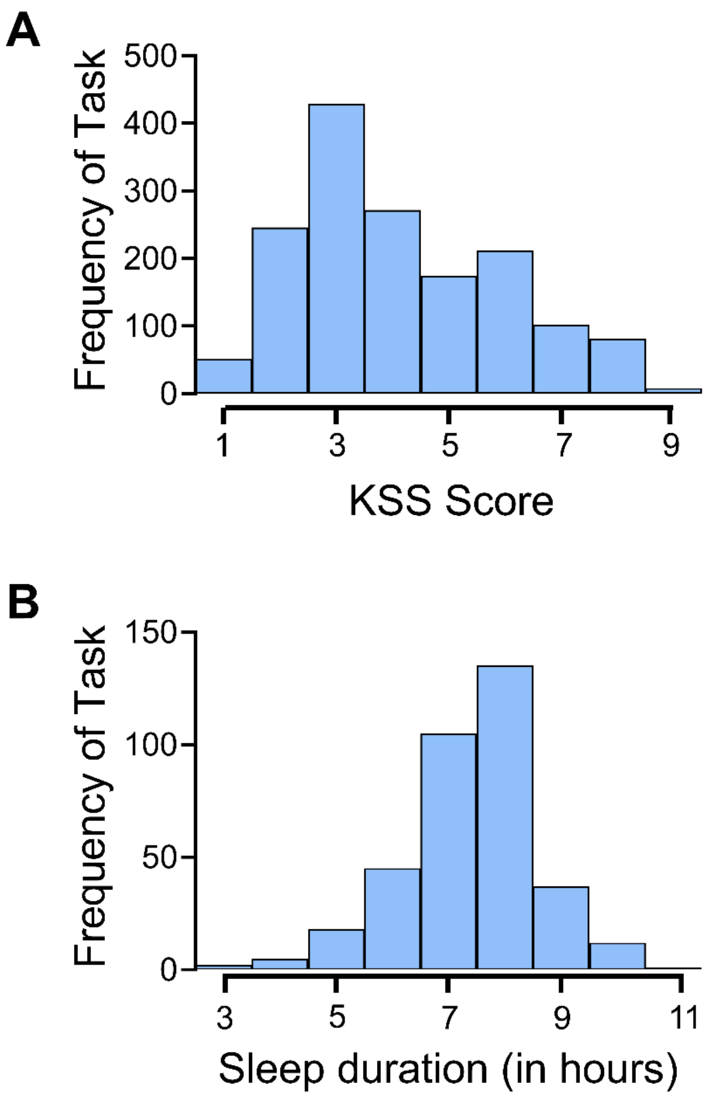


**Fig. S3. Distributions of self-reported mood reports in relation to time of day and sleep-related variables (n=44).** After excluding observations from participants with missing time-awake or sleep-duration data, 44 participants remained. Histograms show the number of completed mood assessments as a function of (A) Karolinska Sleepiness Scale (KSS) score, reflecting participants' reported sleepiness and (B) sleep duration.

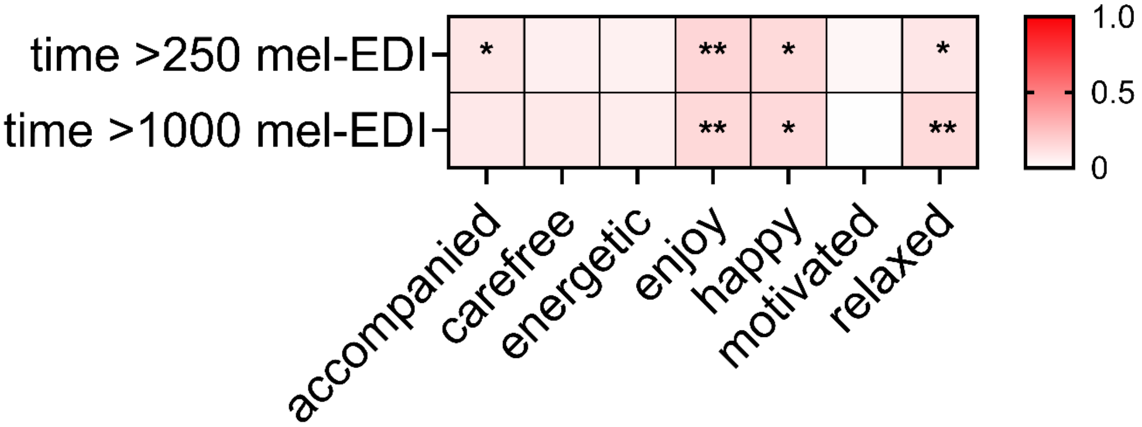


**Fig. S4. Non-covariate-adjusted daily effects of light exposure on self-reported mood (n=48)**. Mood states assessed across panels were feeling accompanied, carefree, energetic, enjoyment, happiness, motivation, and relaxation. Associations were estimated using linear mixed-effects models. Heatmaps display standardised effect sizes, indicating the strength and direction of each association (red, positive; white, negative). Asterisks denote statistically significant associations based on Type III ANOVA: $P < 0.05$ (*), $P < 0.01$ (**), and $P < 0.001$ **(*****). Positive coefficients indicate more positive mood states.

### Tables

**Table S1. Participant demographic and lifestyle characteristics.** Demographic, employment, physical activity, and lifestyle characteristics of participants (n=51).

| | *n* = | Percentage (%) | Mean | Standard Deviation |
|---|---|---|---|---|
| Age | | | 27.35 | 7.03 |
| Gender | | | | |
| Male | 25 | 49.0 | | |
| Female | 26 | 51.0 | | |
| Employment Status | | | | |
| Full time | 31 | 60.8 | | |
| Part time | 0 | 0 | | |
| Student | 15 | 29.4 | | |
| Student + Part time job | 4 | 7.8 | | |
| Unemployed | 1 | 2.0 | | |
| Highest Education | | | | |
| GCSE | 1 | 2.0 | | |
| A Level | 4 | 7.8 | | |
| BSc/BA | 14 | 27.4 | | |
| MSc/MA | 19 | 37.3 | | |
| PhD | 13 | 25.5 | | |
| Alcohol Intake | | | | |
| Never | 2 | 3.9 | | |
| Special Occasions | 11 | 21.6 | | |
| 1-2 times a week | 31 | 60.7 | | |
| 3-4 times a week | 6 | 11.8 | | |
| Daily/almost daily | 1 | 2.0 | | |
| Smoking intake | | | | |
| Smoker | 9 | 17.6 | | |
| Between 1-5 a day | 4 | | | |
| Between 5-10 a day | 2 | | | |
| 10+ a day | 3 | | | |
| Non-smoker | 42 | 82.4 | | |
| Caffeine in a day | | | | |
| 0 cups/cans | 7 | 13.7 | | |
| 1-4 cups/cans | 35 | 68.2 | | |
| 5-7 cups/cans | 6 | 11.8 | | |
| 7+ cups/cans | 3 | 5.9 | | |

| | | |
|---|---|---|
| Exercise | | |
| Never | 3 | 5.9 |
| Rarely | 2 | 3.9 |
| Once a week | 3 | 5.9 |
| A few days a week | 15 | 29.4 |
| Most days | 22 | 43.1 |
| Every day | 6 | 11.8 |

**Table S2. Covariates included in analyses of light exposure and self-reported mood.** Linear mixed-effects model standardised β coefficients and associated Type III ANOVA *P* values for covariates controlled for when assessing associations between light exposure and self-reported mood scores are reported for the full study sample (*n* = 50).

| | Accompanied | Carefree | Energetic | Enjoyment | Happy | Motivated | Relaxed |
|---|---|---|---|---|---|---|---|
| KSS_morning | | | | | | | |
| Standardised β | -0.04 | -0.10 | -0.23 | -0.12 | -0.16 | -0.17 | -0.12 |
| P Value | 0.22 | 0.009 (**) | P<0.001 (***) | P<0.001 (***) | P<0.001 (***) | P<0.001 (***) | 0.002 (**) |
| Sin | | | | | | | |
| Standardised β | -0.006 | -0.02 | -0.16 | -0.09 | -0.06 | -0.07 | -0.02 |
| P Value | 0.74 | 0.32 | P<0.001 (***) | 0.005(**) | 0.04(*) | 0.07(*) | 0.31 |
| Cos | | | | | | | |
| Standardised β | -0.01 | 0.01 | -0.24 | 0.009 | -0.004 | -0.24 | 0.06 |
| P Value | 0.50 | 0.48 | P<0.001 (***) | 0.73 | 0.89 | P<0.05(*) | 0.03(*) |
| Time Awake | | | | | | | |
| Standardised β | 0.05 | 0.04 | 0.73 | 0.27 | 0.24 | 0.57 | -0.03 |
| P Value | 0.70 | 0.56 | P<0.001 (***) | 0.02(*) | 0.008 (**) | P<0.001 (***) | 0.94 |
| I(timeawake^2) | | | | | | | |
| Standardised β | -0.04 | -0.02 | -0.85 | -0.19 | -0.19 | -0.76 | 0.07 |
| P Value | 0.75 | 0.77 | P<0.001 (***) | 0.83 | 0.03 (*) | P<0.05(*) | 0.84 |
| Step_30_mean | | | | | | | |
| Standardised β | 0.02 | 0.01 | 0.06 | 0.007 | 0.003 | 0.07 | -0.02 |
| P Value | 0.29 | 0.52 | 0.02(*) | 0.83 | 0.88 | 0.01(**) | 0.41 |
| Weekdays | | | | | | | |
| Standardised β | -0.05 | -0.06 | -0.05 | -0.11 | -0.10 | -0.005 | -0.08 |

| | | | | | | | |
|---|---|---|---|---|---|---|---|
| P Value | 0.002(**) | P<0.001 (***) | 0.03(*) | P<0.001 (***) | P<0.001 (***) | 0.82 | P<0.001 (***) |
| Age | | | | | | | |
| Standardised β | -0.26 | -0.08 | -0.03 | -0.01 | -0.02 | 0.05 | -0.13 |
| P Value | 0.60 | 0.59 | 0.55 | 0.86 | 0.78 | 0.40 | 0.36 |
| SexFemale | | | | | | | |
| Standardised β | 0.26 | 0.05 | -0.01 | 0.09 | 0.04 | -0.03 | 0.07 |
| P Value | 0.02(*) | 0.67 | 0.87 | 0.29 | 0.68 | 0.68 | 0.46 |
| MCTQ-MSFsc | | | | | | | |
| Standardised β | -0.10 | -0.12 | -0.03 | -0.50 | 0.006 | -0.14 | 0.10 |
| P Value | 0.42 | 0.27 | 0.70 | 0.60 | 0.95 | 0.92 | 0.28 |
| Photoperiod | | | | | | | |
| Standardised β | 0.21 | 0.14 | 0.17 | 0.09 | 0.12 | 0.06 | 0.21 |
| P Value | 0.19 | 0.27 | 0.02(*) | 0.38 | 0.18 | 0.43 | 0.05(*) |

Note. † p < .10. *p* < .05 (*), *p* < .01 (**), *p* < .001 (***). KSS_morning: Karolinska Sleepiness Scale when participant first wakes up; I(timeawake^2): Quadratic effect of time awake; Step_30_mean: amount of steps in 30 minutes before completing questionnaire; MCTQ-MSFsc: Chronotype measured by Munich Chronotype Questionnaire.

**Table S3. Associations between self-reported mood and affective bias within questionnaire sessions.** Pearson's correlation *R* coefficients and associated *P* values are reported for relationships between self-reported mood ratings and affective bias measures collected during the same questionnaire session (*n* = 47).

| | Accompanied | Carefree | Energetic | Enjoyment | Happy | Motivated | Relaxed |
|---|---|---|---|---|---|---|---|
| ECAT_neg_bias | | | | | | | |
| *R* | -0.07 | -0.14 | -0.13 | -0.15 | -0.13 | -0.16 | -0.07 |
| P value | 0.26 | 0.02(*) | 0.03(*) | 0.009(**) | 0.02(*) | 0.006 (**) | 0.23 |
| EREC_neg_bias | | | | | | | |
| *R* | 0.15 | 0.04 | 0.09 | 0.15 | 0.12 | 0.09 | 0.04 |
| P value | 0.01(*) | 0.45 | 0.12 | 0.01(*) | 0.04(*) | 0.15 | 0.47 |
| EMEM_neg_bias | | | | | | | |
| *R* | 0.03 | 0.04 | 0.02 | 0.07 | 0.05 | 0.04 | 0.09 |
| P value | 0.59 | 0.48 | 0.69 | 0.24 | 0.37 | 0.45 | 0.15 |

Note. † p < .10. *p* < .05 (*), *p* < .01 (**), *p* < .001 (***). ECAT_neg bias: the total negative bias for the emotional categorisation task. EREC_neg bias: the total negative bias for the emotional recall task. EMEM_neg bias: the total negative bias for the emotional memory task.

**Table S4. Associations between self-reported mood rhythmicity and affective bias.** Pearson's correlation *R* coefficients and associated *P* values are reported for relationships between participant-level mood rhythm parameters. Mesor and amplitude derived from cosine model fits and measures of affective bias (*n* = 47).

| | Accompanied | Carefree | Energetic | Enjoyment | Happy | Motivated | Relaxed |
|---|---|---|---|---|---|---|---|
| ECAT_neg_bias | | | | | | | |
| Mesor *R* | -0.07 | -0.21 | -0.37 | -0.31 | -0.30 | -0.36 | -0.19 |
| Amplitude *R* | 0.35 | 0.15 | 0.00 | 0.22 | -0.03 | 0.15 | 0.12 |
| Mesor P value | 0.64 | 0.15 | 0.009 (**) | 0.03(*) | 0.03(*) | 0.01(**) | 0.18 |
| Amplitude P value | 0.01(*) | 0.30 | 0.99 | 0.12 | 0.83 | 0.29 | 0.43 |
| EREC_neg_bias | | | | | | | |
| Mesor *R* | 0.22 | 0.05 | -0.03 | 0.20 | 0.19 | -0.01 | 0.08 |
| Amplitude *R* | -0.15 | 0.15 | -0.03 | 0.11 | 0.07 | 0.14 | 0.03 |
| Mesor P value | 0.14 | 0.74 | 0.86 | 0.17 | 0.20 | 0.93 | 0.57 |
| Amplitude P value | 0.29 | 0.30 | 0.86 | 0.43 | 0.63 | 0.32 | 0.86 |
| EMEM_neg_bias | | | | | | | |
| Mesor *R* | 0.16 | 0.17 | 0.25 | 0.27 | 0.24 | 0.27 | 0.21 |
| Amplitude *R* | -0.08 | 0.07 | 0.08 | -0.04 | 0.01 | 0.17 | 0.07 |
| Mesor P value | 0.27 | 0.25 | 0.08 | 0.06 | 0.09 | 0.05(*) | 0.14 |
| Amplitude P value | 0.58 | 0.65 | 0.60 | 0.79 | 0.96 | 0.23 | 0.61 |

Note. † p < .10. *p* < .05 (*), *p* < .01 (**), *p* < .001 (***)**.** ECAT_neg bias: the total negative bias for the emotional categorisation task. EREC_neg bias: the total negative bias for the emotional recall task. EMEM_neg bias: the total negative bias for the emotional memory task.

**Table S5. Acute light exposure effects on self-reported mood: non-covariate-adjusted analysis.** Linear mixed-effects model standardised β and associated Type III ANOVA *P* values are reported for associations between acute light exposure and self-reported mood across temporal averaging windows of 30, 60, 120, 180 and 240 minutes

| | Accompanied | Carefree | Energetic | Enjoyment | Happy | Motivated | Relaxed |
|---|---|---|---|---|---|---|---|
| Mel30mean | | | | | | | |
| Standardised β | 0.02 | -0.002 | 0.21 | 0.29 | 0.06 | 0.20 | -0.02 |
| P value | 0.33 | 0.92 | P<0.001 (***) | 0.25 | 0.01 (*) | P<0.001 (***) | 0.39 |
| Mel60mean | | | | | | | |
| Standardised β | 0.03 | 0.005 | 0.23 | 0.04 | 0.08 | 0.20 | -0.004 |
| P value | 0.12 | 0.82 | P<0.001 (***) | 0.12 | 0.004 (**) | P<0.001 (***) | 0.88 |
| Mel120mean | | | | | | | |
| Standardised β | 0.04 | 0.009 | 0.23 | 0.07 | 0.1 | 0.20 | 0.002 |
| P value | 0.06 | 0.65 | P<0.001 (***) | 0.03(*) | 0.001 (***) | P<0.001 (***) | 0.94 |
| Mel180mean | | | | | | | |
| Standardised β | 0.05 | 0.01 | 0.23 | 0.09 | 0.11 | 0.19 | 0.01 |
| P value | 0.03(*) | 0.53 | P<0.001 (***) | 0.006 (**) | P<0.001 (***) | P<0.001 (***) | 0.55 |
| Mel240mean | | | | | | | |
| Standardised β | 0.05 | 0.01 | 0.21 | 0.10 | 0.11 | 0.17 | 0.03 |
| P value | 0.02(*) | 0.52 | P<0.001 (***) | 0.002 (**) | P<0.001 (***) | P<0.001 (***) | 0.28 |

(*n* = 48).

Note. † p < .10. *p* < .05 (*), *p* < .01 (**), *p* < .001 (***). Mel30mean: light in 30 minutes before self-reported mood questionnaire, Mel60mean: light in 60 minutes before s self-reported mood questionnaire, Mel120mean: light in 120 minutes before self-reported mood questionnaire, Mel180mean: light in 180 minutes before self-reported mood questionnaire, Mel240mean: light in 240 minutes before self-reported mood questionnaire.

**Table S6. Acute light exposure effects on self-reported mood: covariate-adjusted analysis.** Linear mixed-effects model standardised β coefficients and associated Type III ANOVA *P* values are reported for associations between acute light exposure and self-reported mood across temporal averaging windows of 30, 60, 120, 180 and 240 minutes, after adjustment for relevant covariates (*n* = 47).

| | Accompanied | Carefree | Energetic | Enjoyment | Happy | Motivated | Relaxed |
|---|---|---|---|---|---|---|---|
| Mel30mean | | | | | | | |
| Standardised β | 0.03 | 0.01 | 0.07 | 0.01 | 0.05 | 0.05 | 0.05 |
| P value | 0.24 | 0.86 | 0.05(*) | 0.79 | 0.14 | 0.19 | 0.19 |
| Mel60mean | | | | | | | |
| Standardised β | 0.03 | 0.01 | 0.08 | 0.01 | 0.07 | 0.05 | 0.05 |
| P value | 0.18 | 0.74 | 0.04(*) | 0.76 | 0.05(*) | 0.20 | 0.12 |
| Mel120mean | | | | | | | |
| Standardised β | 0.07 | 0.03 | 0.09 | 0.06 | 0.13 | 0.07 | 0.06 |
| P value | 0.03(*) | 0.31 | 0.06 | 0.15 | 0.003 (**) | 0.11 | 0.10 |
| Mel180mean | | | | | | | |
| Standardised β | 0.10 | 0.04 | 0.12 | 0.09 | 0.18 | 0.11 | 0.07 |
| P value | 0.004(**) | 0.26 | 0.02(*) | 0.04(*) | 0.003 (**) | 0.03(*) | 0.10 |
| Mel240mean | | | | | | | |
| Standardised β | 0.12 | 0.04 | 0.15 | 0.11 | 0.19 | 0.13 | 0.10 |
| P value | P<0.001 (***) | 0.30 | 0.005 (**) | 0.03(*) | P<0.001 (***) | 0.02(*) | 0.02(*) |

Note. † p < .10. *p* < .05 (*), *p* < .01 (**), *p* < .001 (***). Mel30mean: light in 30 minutes before self-reported mood questionnaire, Mel60mean: light in 60 minutes before self-reported mood questionnaire, Mel120mean: light in 120 minutes before self-reported mood questionnaire, Mel180mean: light in 180 minutes before self-reported mood questionnaire, Mel240mean: light in 240 minutes before self-reported mood questionnaire.

**Table S7. Acute light exposure effects on affective bias: non-covariate-adjusted analysis.** Linear mixed-effects model standardised β coefficients and associated Type III ANOVA *P* values are reported for associations between acute light exposure and affective bias across temporal averaging windows of 30, 60, 120, 180 and 240 minutes (*n* = 48)..

| | ECAT_neg_bias | EREC_neg_bias | EMEM_neg_bias |
|---|---|---|---|
| mel30mean | | | |
| Standardised β | 0.06 | 0.01 | -0.005 |
| P value | 0.40 | 0.87 | 0.94 |
| mel60mean | | | |
| Standardised β | 0.08 | 0.02 | -0.0006 |
| P value | 0.24 | 0.82 | 1.00 |
| mel120mean | | | |
| Standardised β | 0.14 | 0.02 | -0.007 |
| P value | 0.04(*) | 0.83 | 0.91 |
| mel180mean | | | |
| Standardised β | 0.11 | 0.00002 | 0.04 |
| P value | 0.10 | 1.00 | 0.56 |
| Mel240mean | | | |
| Standardised β | 0.07 | -0.001 | 0.06 |
| P value | 0.31 | 0.988 | 0.32 |

Note. † p < .10. *p* < .05 (*), *p* < .01 (**), *p* < .001 (***). ECAT_neg bias: the total negative bias for the emotional categorisation task. EREC_neg bias: the total negative bias for the emotional recall task. EMEM_neg bias: the total negative bias for the emotional memory task Mel30mean: light in 30 minutes before self-reported mood questionnaire, Mel60mean: light in 60 minutes before self-reported mood questionnaire, Mel120mean: light in 120 minutes before self-reported mood questionnaire, Mel180mean: light in 180 minutes before self-reported mood questionnaire, Mel240mean: light in 240 minutes before self-reported mood questionnaire.

**Table S8. Daily light exposure effects on self-reported mood: non-covariate-adjusted analysis.** Linear mixed-effects model standardised β coefficients and Type III ANOVA *P* values are reported for associations between daily light exposure-time above 250 lux (TAT250) and 1,000 lux (TAT1000) melanopic equivalent daylight illuminance-and each mood score (*n* = 48).

| | Accompanied | Carefree | Energetic | Enjoy | Happy | Motivated | Relaxed |
|---|---|---|---|---|---|---|---|
| TAT250 | | | | | | | |
| Standardised β | 0.10 | 0.06 | 0.05 | 0.16 | 0.15 | 0.03 | 0.10 |
| P value | 0.03(*) | 0.22 | 0.42 | 0.007 (**) | 0.02 (*) | 0.61 | 0.04(*) |
| TAT1000 | | | | | | | |
| Standardised β | 0.10 | 0.09 | 0.07 | 0.15 | 0.15 | 0.0001 | 0.15 |
| P value | 0.07 | 0.08 | 0.34 | 0.004 (**) | 0.01 (*) | 0.998 | 0.002 (**) |

Note. † p < .10. *p* < .05 (*), *p* < .01 (**), *p* < .001 (***). TAT250: Time >250 melanopic EDI and TAT1000: time >1000 melanopic EDI in minutes compared to self-reported mood scores.

**Table S9. Daily light exposure effects on self-reported mood: covariate-adjusted analysis.** Linear mixed-effects model standardised β coefficients and Type III ANOVA *P* values are reported for associations between daily light exposure-time above 250 lux (TAT250) and 1,000 lux (TAT1000) melanopic equivalent daylight illuminance-and each self-reported mood score ($n = 48$).

| | Accompanied | Carefree | Energetic | Enjoy | Happy | Motivated | Relaxed |
|---|---|---|---|---|---|---|---|
| TAT250 | | | | | | | |
| Standardised β | 0.10 | 0.06 | 0.06 | 0.17 | 0.15 | 0.03 | 0.10 |
| P value | 0.03(*) | 0.19 | 0.41 | 0.002 (**) | 0.01 (*) | 0.61 | 0.02(*) |
| TAT1000 | | | | | | | |
| Standardised β | 0.08 | 0.06 | 0.05 | 0.09 | 0.10 | -0.02 | 0.11 |
| P value | 0.10 | 0.18 | 0.53 | 0.12 | 0.09 | 0.81 | 0.02(*) |

Note. † p < .10. *p* < .05 (*), *p* < .01 (**), *p* < .001 (***). TAT250: Time >250 melanopic EDI and TAT1000: time >1000 melanopic EDI in minutes compared to self-reported mood scores.

**Table S10. Participant study schedule.** Schedule provided to participants specifying when each study task should be completed to support adherence to the study protocol.

| Day of study | Tasks to complete | When? |
|---|---|---|
| Day 1: Face-to-face registration meeting | Consent | At baseline visit. Timing arranged with researcher. |
| | Baseline Questionnaire | |
| | Device instructions | |
| Day 2-7 | Sleep questionnaire | When you wake up |
| | Mood questionnaires | 5x a day, e.g. 8am, 11am, 2pm, 5pm, 8pm |
| Day 8: Closing Visit | Sleep questionnaire | When you wake up |
| | Mood questionnaires | 2x during the day |
| | Devices returned. | As arranged |
| +++ Additional days | Sleep questionnaire | When you wake up |
| = 30 minutes a day | Mood questionnaires (5x a day, e.g. 8am, 11am, 2pm, 5pm, 8pm) | 5x a day, e.g. 8am, 11am, 2pm, 5pm, 8pm |

**Dataset S1 (separate file).** Type or paste legend here.
